# Computational ranking-assisted identification of Plexin-B2 in homotypic and heterotypic clustering of circulating tumor cells in breast cancer metastasis

**DOI:** 10.1101/2023.04.10.536233

**Authors:** Emma Schuster, Nurmaa Dashzeveg, Yuzhi Jia, Kibria Golam, Tong Zhang, Andrew Hoffman, Youbin Zhang, Chunlei Zheng, Erika Ramos, Rokana Taftaf, Lamiaa El- Shennawy, David Scholten, Reta B. Kitata, Valery Adorno-Cruz, Carolina Reduzzi, Sabina Spahija, Rong Xu, Kalliopi P. Siziopikou, Leonidas C. Platanias, Ami Shah, William J. Gradishar, Massimo Cristofanilli, Chia-Feng Tsai, Tujin Shi, Huiping Liu

**Affiliations:** Department of Pharmacology, Northwestern University Feinberg School of Medicine, Chicago, IL, USA; Driskill Graduate Program in the Life Sciences, Northwestern University Feinberg School of Medicine, Chicago, IL, USA; Biological Sciences Division, Pacific Northwest National Laboratories, Richland, WA, USA; Department of Biomedical Informatics, School of Medicine, Case Western Reserve University, Cleveland, OH, USA; Division of Hematology and Oncology, Department of Medicine, Northwestern University Feinberg School of Medicine, Chicago, IL, USA; Division of Hematology and Medical Oncology, Department of Medicine, Weill Cornell School of Medicine, New York, NY, USA; Department of Pathology, Northwestern University Feinberg School of Medicine, Chicago, IL, USA; Robert H. Lurie Comprehensive Cancer Center, Northwestern University Feinberg School of Medicine, Chicago, IL, USA

## Abstract

Metastasis is the cause of over 90% of all deaths associated with breast cancer, yet the strategies to predict cancer spreading based on primary tumor profiles and therefore prevent metastasis are egregiously limited. As rare precursor cells to metastasis, circulating tumor cells (CTCs) in multicellular clusters in the blood are 20-50 times more likely to produce viable metastasis than single CTCs. However, the molecular mechanisms underlying various CTC clusters, such as homotypic tumor cell clusters and heterotypic tumor-immune cell clusters, are yet to be fully elucidated. Combining machine learning-assisted computational ranking with experimental demonstration to assess cell adhesion candidates, we identified a transmembrane protein Plexin- B2 (PB2) as a new therapeutic target that drives the formation of both homotypic and heterotypic CTC clusters. High PB2 expression in human primary tumors predicts an unfavorable distant metastasis-free survival and is enriched in CTC clusters compared to single CTCs in advanced breast cancers. Loss of PB2 reduces formation of homotypic tumor cell clusters as well as heterotypic tumor-myeloid cell clusters in triple-negative breast cancer. Interactions between PB2 and its ligand Sema4C on tumor cells promote homotypic cluster formation, and PB2 binding with Sema4A on myeloid cells (monocytes) drives heterotypic CTC cluster formation, suggesting that metastasizing tumor cells hijack the PB2/Sema family axis to promote lung metastasis in breast cancer. Additionally, using a global proteomic analysis, we identified novel downstream effectors of the PB2 pathway associated with cancer stemness, cell cycling, and tumor cell clustering in breast cancer. Thus, PB2 is a novel therapeutic target for preventing new metastasis.

## Highlights

Computational ranking-assisted identification of candidates associated with metastasis and patient outcomes.

Plexin-B2 on tumor cells promotes CTC clustering and lung metastasis of breast cancer.

Plexin-B2 interactions with Sema4C on tumor cells and Sema4A on monocytes drive tumor cluster formation and enhance lung metastasis *in vivo*.

## Introduction

Rapid development of cutting-edge machine learning and deep learning has facilitated the accumulation and transformation of bioinformatic data into valuable knowledge and new discoveries^1–5^. Reciprocally, comprehensive experimental determination solidifies computational analysis-based data mining and prediction. Our study seeks to synergize the power of bioinformatic analysis and experimental exploration for cancer discoveries. Despite great progress in cancer prevention and medical care, 1 woman out of 8 in the United States is still predicted to develop breast cancer in her lifetime ^6^, with expression of estrogen receptor (ER), progesterone receptor, epidermal growth factor receptor 2 (HER2), or triple-negative for all three ^7, 8^. Most Stage IV breast cancers are diagnosed as triple-negative breast cancer (TNBC) with a median survival of only 8.8 months after metastasis to the lungs, brain, and liver ^9, 10^. As such, our goal is to collect and mine phenotype-associated proteomic data for computational ranking of candidates, and therefore identify viable therapeutic targets for breast cancer metastasis.

Cancer is disseminated by circulating tumor cells (CTCs) that shed off the primary tumor and are capable of seeding and regenerating tumors in distant organs, including lung, liver, brain, and bone, due to their inherent and acquired properties, such as stemness and/or proliferation^11–14^. The dogma of single CTC-mediated cancer dissemination has been challenged by the detection of rare CTC clusters in the blood of patients with advanced breast cancer^11, 15–19^. The existence of CTC clusters predicts unfavorable outcomes^11, 15–19^, as they are more likely to seed metastases than single CTCs^11, 20–24^. Due to the devastating features of metastatic disease, it is imperative to discover the diverse mechanisms of CTC clusters in cancer.

CTC clusters may contain multiple cell types. Homotypic tumor cell clusters are often enriched with tumor cells that have stem cell properties^16, 25–27^, whereas heterotypic clusters contain mixtures of tumor and immune cells in which the tumor cells can evade immune cell attack and show proliferation advantages^19, 28^. Myeloid cells (neutrophils and monocytes) account for a large proportion of immune cells that interact with CTC clusters^19^. Two cellular mechanisms have been proposed for CTC cluster formation^18^; one is collective dissemination or cohesive shedding^11, 29^, and another is tumor cell aggregation^16, 25^, which would lead to both homotypic and heterotypic tumor clusters. While both homotypic and heterotypic CTC clusters can promote metastasis in association with unfavorable overall survival (OS)^18^, the molecular mechanisms have yet to be fully elucidated. Our previous work and that of others have identified a few molecular drivers of homotypic tumor cell clusters, including tumor-initiating cell markers and receptors^16, 25, 26^ and molecules in cell junctions or adhesive structures^11^. Integrins on neutrophils may enhance cell cycle progression in heterotypic CTC clusters^18, 19^.

It remains an open question whether primary tumor profiles can be used to predict such drivers for early intervention, therefore intervene and prevent homotypic and heterotypic tumor cluster formation. We hypothesize that aggregation or cohesion phenotype-related adhesion proteins can regulate CTC cluster formation. Taking advantage of systems biology and mass spectrometry (MS)-based global proteomic data, we developed a novel computational ranking to assess all cell adhesion molecule candidates in breast cancer. As proof-of-concept, our studies revealed a transmembrane protein, Plexin-B2 (PB2), as the top candidate that is up-regulated in primary tumors and enhances the formation of both homotypic tumor cell clusters and heterotypic CTC- myeloid cell clusters in metastasis of breast cancer, especially TNBC.

PB2 is a single-pass transmembrane plexin family member, and its primary function is to direct neural cell growth and migration in brain development; it also plays a role in the function of the vascular and endocrine systems, wound healing, monocyte function, and stem cell dynamics in human embryonic stem cells and neuroprogenitor cells^30–37^. However, PB2’s molecular mechanism has not previously been studied in the context of CTC clustering and metastatic breast cancer.

## Results

### Computational ranking of cell adhesion molecules in breast cancer

To seek an unbiased *in silico* screen of candidate proteins that may regulate CTC cluster formation, we first ranked all cell adhesion molecules (N=608, derived from the Molecular Signature Database^38–40^ of the Gene Ontology Biological Processes^41, 42^) according to their relative protein abundance in human breast tumors and TNBC cells, in TNBC tumor tissue versus normal adjacent tissues, and in cancer-derived extracellular vesicles (EVs) (**Figure 1A**). By analyzing the tandem MS proteomic datasets of 122 treatment-naive human breast tumors^43^, 398 adhesion proteins were detected and ranked based on the number of spectra per protein (**Figure 1A**, **Figure S1A, Supplementary Excel 1-tab 1**). To further determine the relevance of adhesion proteins in cancer cells, we examined the MS profiles of human TNBC cells^44^ in which 252 expressed adhesion proteins were ranked based on peptide-spectrum match (PSM) counts (**Figure 1A**, **Figure S1B, Supplementary Excel 1-tab 2**). The overlapped list includes previously characterized surface protein regulators of homotypic CTC clusters, CD44^16^, ICAM1^25^, and CD81^26^ within the top 50-100 most abundant proteins on cancer cells (**Figure S1A–B**), validating the potential relevance of listed adhesion molecules. Moreover, we performed sensitive proteomic analyses of TNBC human tumor tissue voxels and adjacent normal mammary tissue voxels dissected by lase capture microdissection, identifying 627 differentially expressed proteins in tumor tissues (**Figure 1A**, **Supplementary Excel 1-tab 3**).

**Figure 1.**
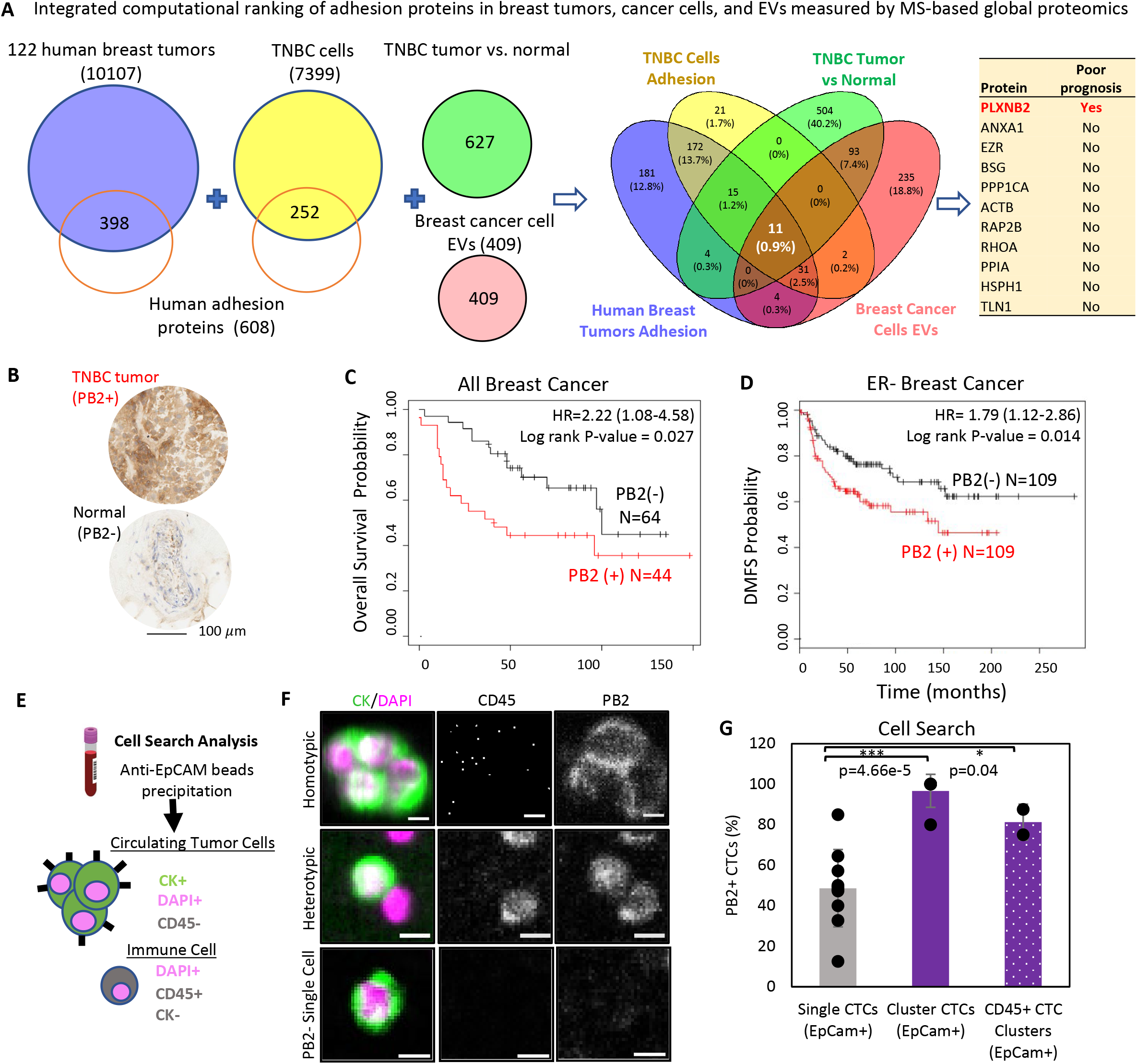
PB2 expression in breast tumors is associated with poor prognosis and enriched in CTC clusters. **A**) Schematic of integrating computational rankings of adhesion proteins into a four-way Venn diagram, based on mass spectrometry proteomic abundance in primary breast tumors (N=122) and TNBC cell lines, altered protein expression in TNBC tumor vs. normal tissue voxels, and breast cancer cell-secreted EVs; to identify novel candidates associated with patient outcomes. **B**) Representative IHC images of human PB2^+^ TNBC tumor and PB2^-^ normal breast tissue (adjacent to tumor) from a patient with TNBC. **C**) KM plot^42^ for overall survival of all patients with breast cancer, divided by best cut-off of high vs. low PB2 protein expression in primary tumors, N=108. **D**) KM plot^42^ for distant metastasis-free survival (DMFS) of patients with ER^-^ breast cancer, divided by median cut-off of high vs. low PB2 mRNA expression, N=218. **E**) Schematic depicting the patient blood sample analysis workflow for CTC analysis on CellSearch. **F**) Representative CellSearch images of a homotypic PB2^+^ CTC-CTC (CD45^-^CK^+^DAPI^+^) cluster, a heterotypic PB2^+^ CTC-WBC (CD45^+^CK^-^DAPI^+^) cluster, and a single PB2^-^ CTC, stained after anti-EpCAM bead enrichment. Scale bar = 5 µm. **G**) Portion of CellSearch-analyzed PB2^+^ CTCs (%) in single CTCs, homotypic CTC clusters, and heterotypic CTC-WBC clusters, as shown in (F), N=10. Data are presented as mean values +/- SD, p-values reported are from two-sided unpaired t-tests.

To further assess the relevance of candidate proteins contributing to the microenvironment for metastasis^45–48^, we performed MS proteomic analyses of small EVs secreted by breast cancer cells (MCF7, SKBR3, BT4T4, and MDA-MB-231) and normal immortalized epithelial cells (MCF10A and MCF12A). Most of the EV proteins were glycoproteins and membrane-bound proteins, and many were differentially enriched in cancer EVs (**Figure S1C–D, Supplementary Excel 1-tabs 3-4**). Across the four MS datasets of human breast tumors, TNBC cells, TNBC tumor vs. normal tissue, and cancer EVs, we created a four-way Venn diagram and identified 11 overlapped proteins (**Figure 1A**, **Figure S1D**). Among the 11 proteins, the top candidate was PB2, which had a high abundance rank in breast tumors and TNBC cells, was highly up-regulated both in tumor tissues and cancer EVs versus their normal controls, and was the only protein associated with an unfavorable prognosis in breast cancer (**Figure 1A**, **Figure S1A–E, Supplementary Excel 1-tabs 5-6**).

### PB2 expression is associated with unfavorable survival and enriched in CTC clusters

In KMPlotter^49^ analysis of human breast cancer datasets^50^, we found that high PB2 protein expression is negatively associated with OS in all breast cancers and with distant metastasis-free survival (DMFS) in ER-negative breast cancers or in grade 3 breast cancers (**Figure 1B–D**, **Figure S2A**). We also analyzed the PB2 protein expression in a tissue microarray of advanced breast cancers collected at Northwestern University and observed high PB2 expression across different breast cancer subtypes (**Figure S2B–D**). Notably, most patients with PB2^high^ breast cancer had detectable blood CTCs, stained as cytokeratin (CK)^+^DAPI^+^CD45^-^ cells after being enriched with anti-EpCAM magnetic beads in an FDA-approved CellSearch® analysis (**Figure 1E–F**, **Figure S2E**).

We then analyzed the expression of PB2 in CTCs of the patients with stage III-IV breast cancer using two previously established methods^16, 25^: CellSearch with blood collected into fixative-containing CellSave tubes (N=10), and live cell flow cytometry^18, 25^ (**Figure S3A–B**) with blood drawn into EDTA tubes (N=17). PB2 was analyzed in CTCs in addition to other channels for epithelial markers CK and/or EpCAM, white blood cell (WBC) or leukocyte marker CD45, and DAPI (nuclear DNA on CellSearch or negative viability for flow cytometry). Compared to single CTCs, PB2 expression was significantly higher in CTC clusters, both homotypic CTC-CTCs and heterotypic CTC-WBC clusters, as analyzed on CellSearch (**Figure 1F–G**). Flow cytometry analysis confirmed the CellSearch® results that showed a significantly larger proportion of CTC clusters were PB2-positive than single CTCs (**Figure S3B–E**).

### PB2 promotes CTC clustering and stemness

After identifying PB2 enrichment in CTC clusters, we continued to determine its function in tumor cell clustering in specific breast cancer subtypes such as TNBC and HER2^+^ in which metastasis is common. To do this, we sorted luciferase 2-eGFP (L2G)-labeled PB2^high^ and PB2^low/-^ tumor cells from our established TNBC patient-derived xenograft (PDX) models (TN3) which develop spontaneous lung micrometastases after orthotopic implantation at mouse mammary fat pads^13^ (**Figure 2A**). Next, the clustering of primary PDX tumor cells on collagen-coated plates was monitored over time using the IncuCyte Live Cell Imager® as previously described^16^. After 6-8 h of clustering, PB2^high^ cells formed significantly more clusters (>2-3 cells) than PB2^low^ cells (**Figure 2B–C**). We then knocked down PB2 in multiple human and mouse TNBC cell lines, MDA-MB-231, 4T1, and HS578T, as well as HER2^+^ cell line SKBR3, using both SmartPool siRNA (siPB2) and individual siRNAs (siPB2-09, -10, -11). The reduction or loss of the full-length (∼250 kDa) and truncated (∼75 kDa) PB2 was shown by immunoblotting (**Figure S4A**) and flow cytometry (**Figure S4B–K**). We observed that cells transfected with siPB2 started to show a slower growth (confluence) than the control cells at 24-96 h after seeding, whereas siPB2-11 with a lower knockdown (KD) efficiency did not have a significant effect on the growth (**Figure S4A**). When clustering was monitored over a short period of 4-6 h with minimal influence by cell proliferation/growth, loss of PB2 significantly reduced the cluster size of all four tested models, including MDA-MB-231 and MDA-MB-468 cells (**Figure 2D–G**) as well as mouse (4T1) and human HS578T lines (**Figure S4F–K**), suggesting that PB2 is a general promoter for CTC clustering in breast cancers.

**Figure 2.**
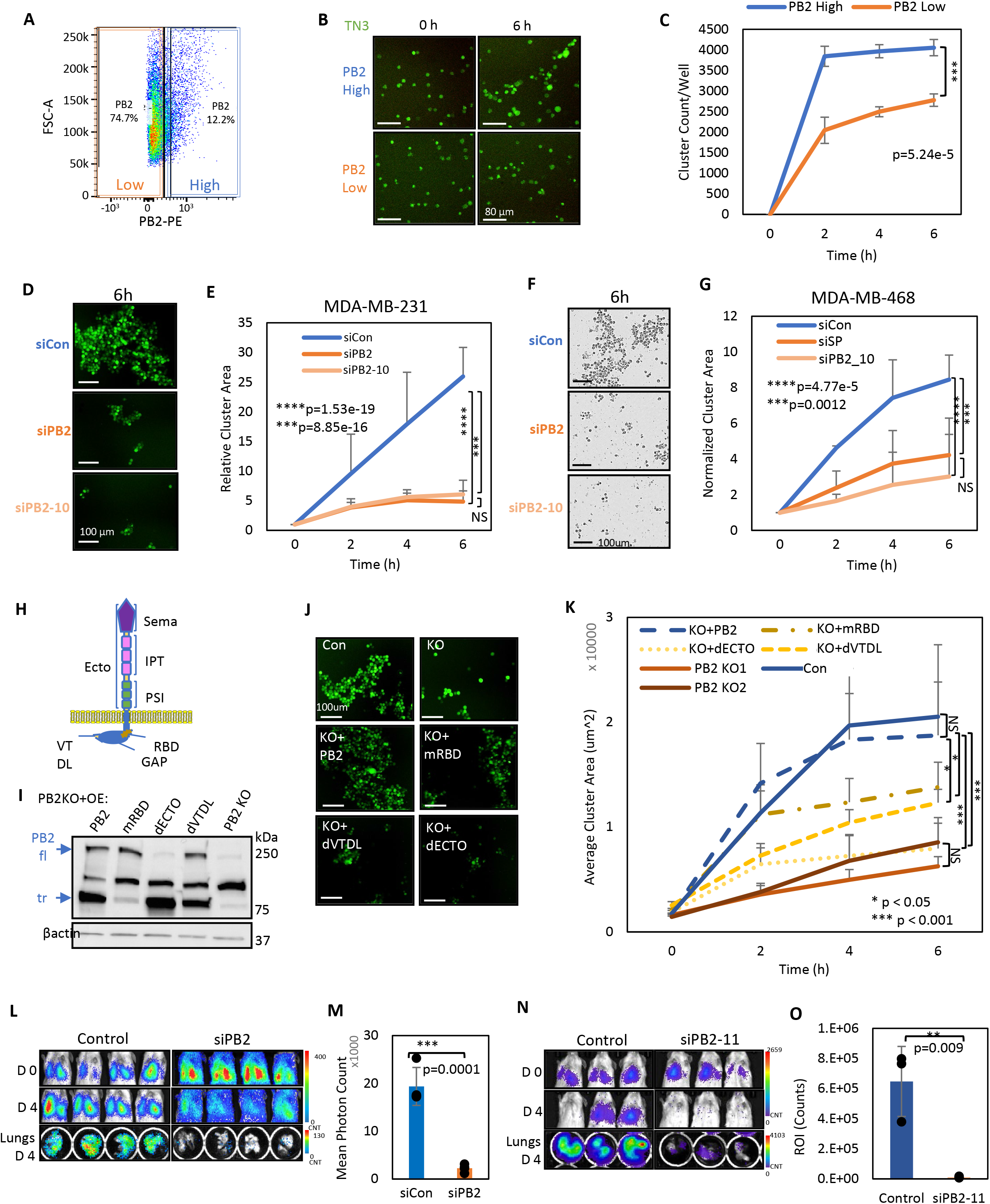
PB2 promotes CTC clustering and lung colonization in TNBC. **A)** Flow panel showing the gating of PB2^+^ and PB2^-^ cells sorted from dissociated primary TN3 L2G^+^ PDX cells for clustering in (B-C). **B & C)** Representative images at 0 and 6 h of clustering (B) and cluster count curves of sorted PB2^high^ and PB2^low^ TN3 PDX cells (C), measured by IncuCyte imaging system; N=3 experiments with at least 5 technical replicates each. **D & E)** Representative images (D) and cluster area growth curves (E) of MDA-MB-231 cells treated with control (siCon), PB2 KD via SmartPool siRNA (siPB2), and single target siRNA (siPB2-10), N=3 experiments with at least 5 technical replicates each. **F & G)** Representative images (F) and cluster area growth curves (G) of MDA-MB-468 cells treated with siCon, PB2 KD via siPB2 SmartPool (siPB2), and siPB2-10. **H**) Schematic of PB2 domains: Extracellular (Ecto) domains include **Sema** = Semaphorin domain; **IPT** = Ig-like fold domain; and **PSI** = Plexin-semaphorin-integrin domain. Intracellular domains include **RBD** = Rho-binding domain; **GAP** = GTPase activating protein domain; and **VTDL** = PDZ-domain binding site (Rho-GEF binding). **I**) Immunoblots of overexpressed PB2 mutants with either full-length (fl) or truncated (tr) depletions: mutated RBD (mRBD) tr, depleted Ecto domain (dECTO) fl and depleted VTDL (dVTDL) tr in MDA-MB-231 PB2 KO cells. **J & K)** Representative images (J) and cluster area growth curves (K) of MDA-MB-231 PB2 KO clusters with overexpression of PB2 full-length or mutants (mRBD, dECTO, or dVTDL) over 6 h as measured via IncuCyte; Con vs. KO+PB2 p=0.67, Con vs. KO1 p=0.0002, Con vs. KO2 p=1.43e-6, KO+PB2 vs. KO+mRBD p=0.04, KO+PB2 vs. KO+dECTO p=1.69e-6, KO+PB2 vs. KO+dVTDL p=0.005, KO+PB2 vs. KO1 p=0.0002. **L-O)** IVIS bioluminescence images (L, N) of NSG mice on days 0 and 4, and quantified signals of metastasis (M, O) of dissected lungs *ex vivo* on day 4 after tail vein injection of MDA-MB- 231 cells transfected with siCon, siPB2 (L), or siPB-11 (N), N=4 (L-M) and 3 (N-O) biological replicates. Data are presented as mean values +/- SD, p-values reported are from two-sided unpaired t-tests.

To further complement the temporary gene KD experiments, we next generated multiple pooled populations of PB2 knockout cells (KO1 and KO2) using CRISPR-Cas9 and gRNAs (**Figure S5A–D**). Possibly due to selective pressure, PB2 KO cells did not slow down the growth or proliferation as the KD cells had (**Figure S5E**), but still caused a similar reduction in tumor cell clustering (**Figure S5F–G**), demonstrating that PB2 is necessary for tumor cell clustering and acts independently of the proliferation phenotype.

PB2 is a large single-pass transmembrane protein made up of multiple domains (**Figure 2H**): the extracellular region (ECTO) including a semaphorin domain (Sema) responsible for binding to receptor/ligands, three Ig-like fold domains and three plexin-semaphorin-integrin domains; a transmembrane domain; the intracellular Rho-Binding Domain (RBD); the intracellular GTPase activating protein domain, and the intracellular PDZ-binding domain (VTDL). Using the PB2 mutant constructs originally designed by the Friedel Lab^51^ (deposited with Addgene), we overexpressed the full-length PB2 and its various mutant forms (dECTO, mRBD, and dVTDL) back into PB2 KO cells and validated the expression using immunoblotting (**Figure 2I**) or flow cytometry (**Figure S5H**). While full-length PB2 overexpression effectively rescued the clustering of PB2 KO cells, the PB2 mutant depleting the ECTO domain (dECTO) had the lowest cluster- rescuing function, followed by the mutants depleting or modifying the intracellular signaling domains (dVTDL and mRBD) (**Figure 2J–K**). These data demonstrate that both extracellular and intracellular signaling properties in full-length PB2 are required to promote tumor cell clustering.

As stemness is often required for CTCs to mediate metastasis with properties of self-renewal, plasticity, and tumorigenicity^13, 52–54^, we next determined if PB2 regulates stemness-related mammosphere formation of breast cancer cells *in vitro*^16^. We found that KD or KO of PB2 significantly reduced the number of mammospheres formed in TNBC cells (**Figure S5I–L**). The mammosphere phenotype in PB2 KO cells was restored by overexpression of the full-length PB2 and to a lesser extent by overexpressed dECTO PB2 mutant, suggesting that the extracellular domain of PB2 is necessary for its functions in promoting self-renewal of tumor cells (**Figure S5K–L**).

Given that CTC clusters are powerful precursors to metastasis, we wanted to determine if PB2 levels would impact CTC dissemination to the lungs. Within 4 days after tail vein injection for experimental metastasis, the mice injected with L2G-labeled^13^ MDA-MB-231 KD cells either by siPB2 or si-PB2-11 had 80-90% less metastasis to the lungs compared to the control group (**Figure 2L–O**), possibly independent of the siPB2 effects on cell growth or proliferation, which was not significantly affected by siPB2-11 (**Figure S4A**). These data suggest that PB2 promotes CTC clustering, self-renewal, and lung colonization.

### PB2 signals through Sema4C to promote CTC clustering

To elucidate the molecular mechanism by which PB2 facilitates CTC cluster formation, we investigated whether in breast cancer PB2 interacts with any of its canonical ligands on the cell surface, such as single-pass transmembrane proteins, semaphorin (Sema) family members Sema4A, 4C, 4D, and 4G^51, 55–60^. From our ranked list of adhesion proteins, we found expression of Sema4C, Sema4D, and Sema4A in treatment-naïve breast cancer^43^ (**Figure S6A**); however, only Sema4C was dramatically detected in breast cancer cells (human/mouse cell lines and/or PDX tumor cells) via immunoblot analyses and mass spectrometry (**Figure 3A–B**, **Supplementary Excel 1-tab 7**). Based on publicly available databases, we also found that high Sema4C mRNA expression in breast cancer correlates with unfavorable DMFS and OS (**Figure S6B–C**). We hypothesized that Sema4C is a primary ligand of PB2 in driving homotypic CTC-CTC clusters in metastasis.

**Figure 3:**
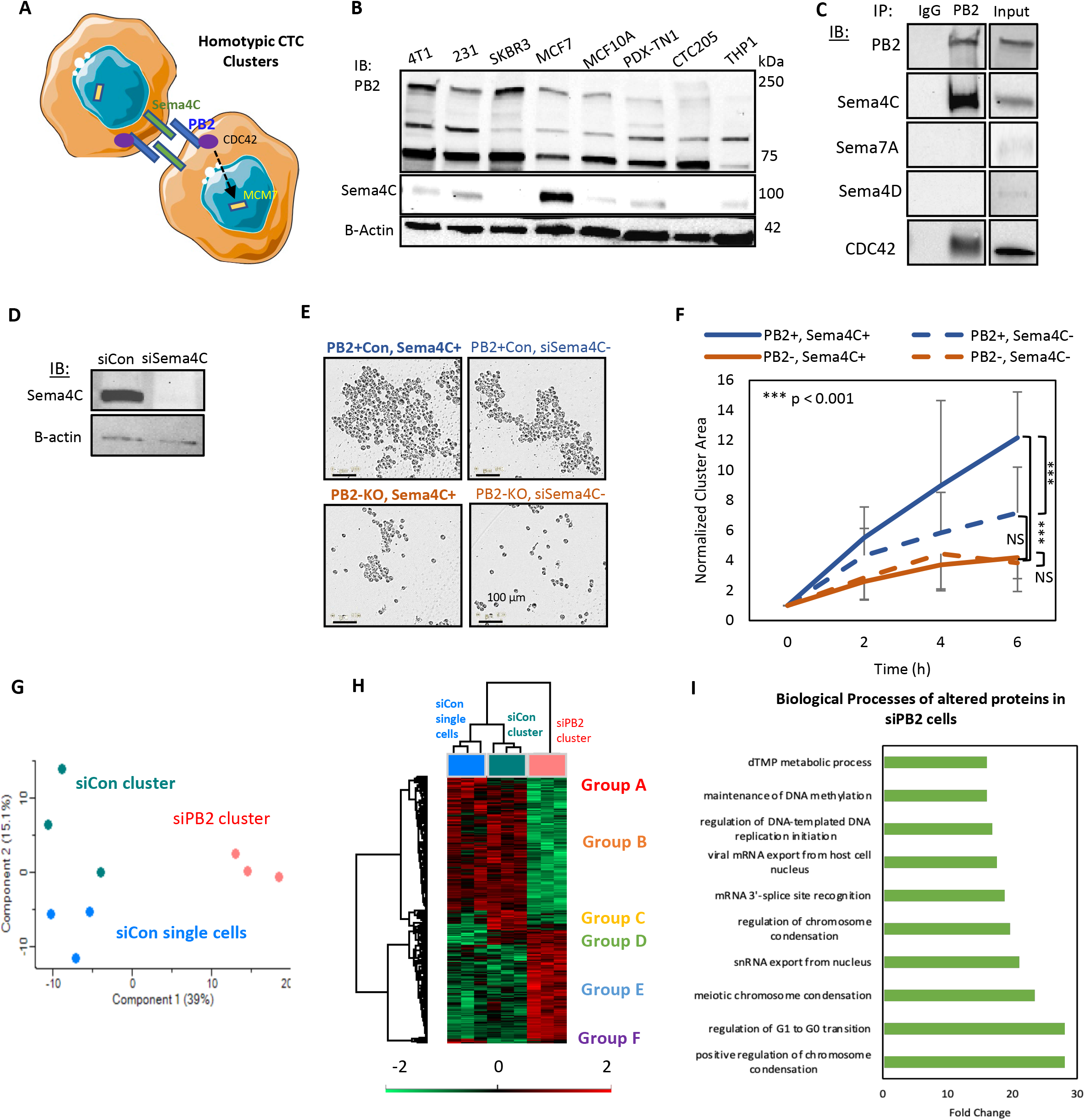
PB2 interacts with Sema4C to promote homotypic clustering of TNBC cells. **A**) Schematic of PB2-Sema4C intercellular interactions in homotypic tumor cell clustering between two cancer cells. **B**) Immunoblots of PB2 and Sema4C expression in different breast cancer cell lines and PDX models using antibodies that are specific to detect both human and mouse isoforms. **C**) Immunoblots of Sema4C, CDC42, and PB2 in the protein complex immunoprecipitated by anti-PB2 antibody compared to IgG. **D**) Immunoblot image with reduced Sema4C after Sema4C KD in MDA-MB-231 tumor cells. **E & F)** Representative images (E) and average cluster growth area curves (F) of MDA-MB-231 control and PB2 KO cells in co-culture with Con or Sema4C KD cells, measured by IncuCyte, N=3 experiments with at least 5 technical replicates each, PB2^+^Sema4C^+^ vs. PB2^+^Sema4^-^ p=1.86e-9, PB2^+^Sema4C^+^ vs. PB2^-^Sema4C^-^ p=6.81e-10, PB2^-^Sema4C^+^ vs. PB2^-^Sema4C^-^ p=0.22, PB2^+^Sema4C^-^ vs. PB2^-^Sema4C^-^ p=0.47. **G**) Principal component analysis clusters of MDA-MB-231 single cells, siCon clusters, and siPB2 clusters analyzed from global mass spectrometry with N=3 replicates per group. **H**) Heat map of differentially expressed proteins from global mass spectrometry analysis of siCon single cells (blue) N=3, siCon clusters (green) N=3, and siPB2 clusters (red) N=3 **I**) Gene ontology biological processes analysis of significantly up-regulated and down-regulated proteins in PB2 control vs. PB2 KD clusters in breast cancer cells analyzed by mass spectrometry. Data are presented as mean values +/- SD, p-values reported are from two-sided unpaired t-tests.

To confirm if Sema molecules interact with PB2 in breast cancer cells, we performed a co-immunoprecipitation for human PB2 using clustered TNBC cells; and found that Sema4C was specifically enriched in the PB2 protein complex, whereas Sema4D and Sema7A were nearly undetectable (**Figure 3C**), demonstrating Sema4C-PB2 interactions in the homotypic CTC clusters. We also detected CDC42 in the anti-PB2 pull-down protein complex (**Figure 3C**). CDC42 is a Rho GTPase family member known to be regulated by PB2 signaling and implicated in migration of cancer cells^61, 62^, indicating the involvement of PB2 signaling in CTC clusters (**Figure 3A**).

To determine the functional importance of Sema4C in CTC clustering, we knocked down Sema4C alone or together with PB2 KO in TNBC cells (**Figure 3D**). Sema4C KD alone mimicked PB2 KO in reducing the average size of tumor cell clusters, whereas loss of both PB2 and Sema4C together did not have an additional impact on cluster size compared to PB2 KO alone (**Figure 3E–F**). This indicates that these two interacting proteins might belong to the same interaction pathway necessary for homotypic CTC clustering.

To gain a better understanding of the downstream molecules and pathways in PB2-mediated CTC clustering, we performed a MS-based global proteomics analysis of MDA-MB-231 breast cancer cells in suspension before and after clustering. Principal component analysis indicates three groups of samples and Heatmap comparisons show 6 distinct groups of protein signatures, up-regulated or down-regulated within 4 h of clustering or by siPB2-mediated KD (**Figure 3G–H**, **Supplementary Excel 2**). Additional GO analysis of the altered proteins in siPB2 cells include biological processes in chromosome condensation, cell cycle, RNA export, DNA methylation, and others (**Figure 3I**), consistent with the phenotypes of stemness, clustering, and cell proliferation. Groups C and F, with clustering-specific up-regulation and down-regulation in protein expression, respectively, identified pathways in protein folding and hydroxylation, UV protection, and response to gamma radiation, all of which were over 10-fold enriched in PB2^+^ tumor cell clusters (**Figure S7A–B**), providing novel insight into how PB2 tunes cellular programming or modifies the post-translational responses to stress from single to clustered cells.

In search of PB2 downstream target proteins related to TNBC phenotypes, we found that MCM7, an essential component of the DNA helicase, was one of the top candidates significantly decreased in the cells with PB2 KD (**Figure S7C**). MCM7 has previously been implicated in cancer cell proliferation and invasion, suggesting that it may help promote cancer stemness and metastasis^63^, thus making it an interesting target for clustering studies. TNBC cells transfected with siMCM7 for KD of MCM7 showed a compromised clustering within 6 h (**Figure S7D–E**). Thus, our data demonstrate a novel mechanism by which PB2 can promote clustering through influencing MCM7.

### PB2-dependent heterotypic clustering signals through Sema4A on monocytes

In addition to MCM7, the global mass spectrometry analysis of PB2 KD cells revealed many biological processes at 10- to 40-fold enrichment, such as positive regulation of the immune response to tumor cells and myeloid cell (astrocyte) activation (**Figure S7A**), indicating that PB2 in tumor cells may impact interactions with immune cells. Since immune cells, especially myeloid cells, have been found in heterotypic CTC clusters driving cell cycle progression, proliferation, and/or immune evasion^19, 64^, we proposed to determine what, if any, role PB2 plays in heterotypic CTC-immune cell clusters (**Figure 4A**).

**Figure 4:**
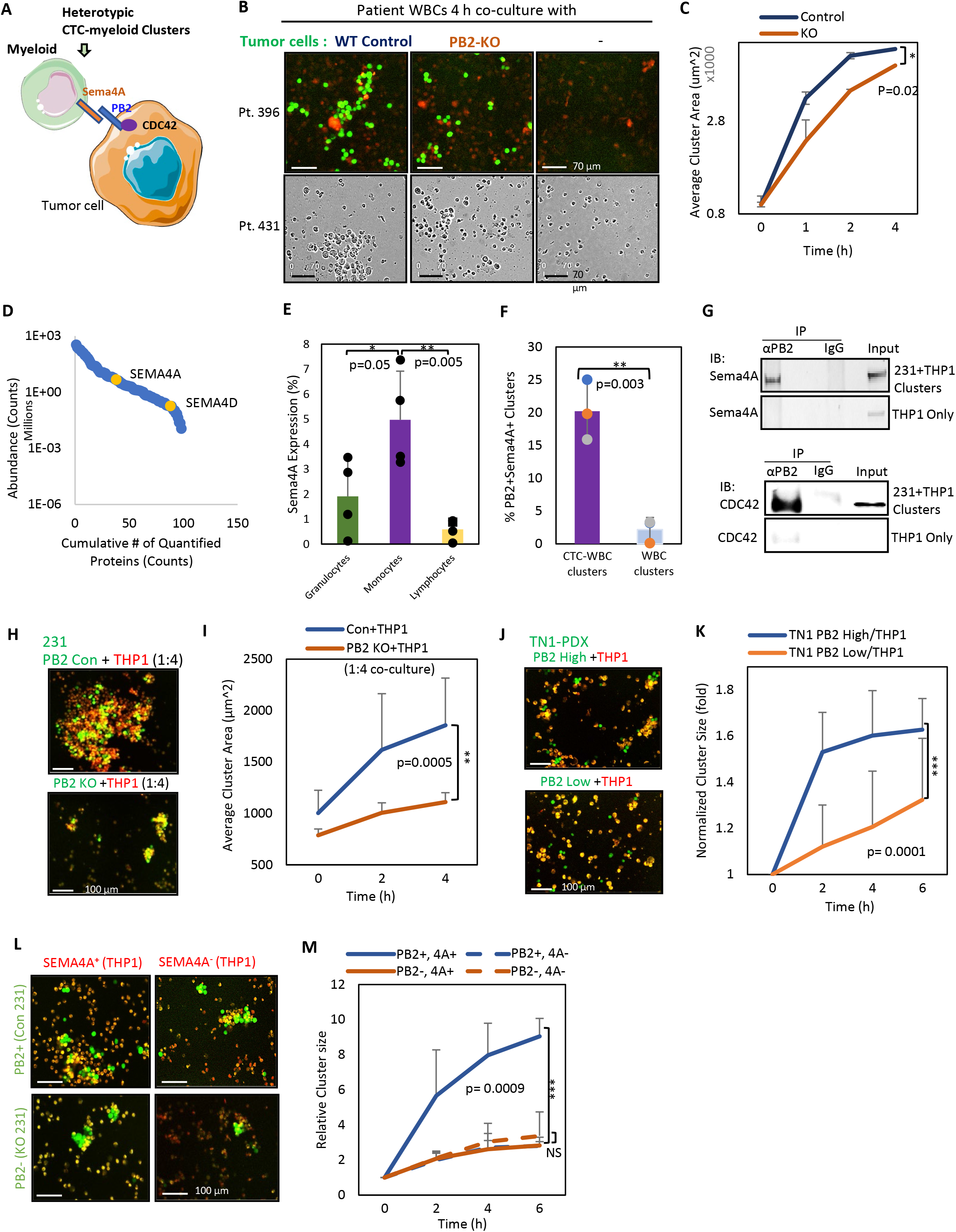
PB2 binding to Sema4A on monocytes promotes heterotypic clustering. **A**) Schematic of the interactions between breast cancer cell PB2 and monocyte Sema4A for heterotypic cluster formation. **B**) Representative images at 4 h co-culture of *ex vivo* heterotypic clusters of L2G^+^ MDA-MB- 231 control or PB2 KO cells with WBCs from breast cancer patients (Pt). Top row: fluorescent images with WBCs from Pt 396 (minimal cytotox red-stained dead cells). Bottom row: brightfield images with WBCs from Pt 431. **C**) Area curves of heterotypic tumor cell clusters co-cultured with WBCs from patients (N=4) as shown in (B), measured by Incucyte imaging. Data are reported as mean +/-SD, with p-values from two-sided unpaired t-tests. **D**) Abundance ranking of 96 identified adhesion/surface proteins in human monocytes (N=3), based on mass spectrometry (MS) quantitative analysis (https://doi.org/10.1038/s41598-020-61356-w). **E**) Sema4A expression on granulocytes, monocytes, and lymphocytes from patients with advanced breast cancer (N=4). **F**) Percent of heterotypic CD45^+^EpCAM^+^ CTC-WBC clusters vs. CD45^+^EpCAM^-^ WBC clusters that express both PB2 and Sema4A, as analyzed by flow analysis, N=4 patients. **G**) PB2 immunoprecipitation of MDA-MB-231 tumor-monocyte (THP1) clusters or THP1 monocytes only, blotted for Sema4A and CDC42. **H & I)** Representative images at 4 h (H) and cluster area curves (I) of heterotypic clustering with MDA-MB-231 tumor cells (green) and THP1 monocytes (red) mixed at a 1:4 ratio, reported as mean +/-SD, N=3 with at least 3 technical replicates each. **J & K)** Representative images at 6 h (J) and cluster size curves (K) of heterotypic clustering with L2G^+^ TN1 PDX tumor cells (green, sorted by PB2^high^ and PB2^low^) and THP1 monocytes (red), N=2 experiments with at least 3 technical replicates each. **L & M)** Representative images at 6 h (L) and cluster size curves (M) of MDA-MB-231 WT (Con) or PB2 KO cells cocultured with THP1 monocytes (sorted by Sema4A+/- expression), N=3 experiments with at least 5 technical replicates each. Data are presented as mean values +/- SD, p-values reported are from two-sided unpaired t-tests.

In co-culture suspension, we mixed PB2 WT or KO MDA-MB-231 cells with patient WBCs. The PB2 WT cells formed larger heterotypic clusters with WBCs than PB2 KO cells without affecting tumor cell viability (**Figure 4B–C**, **Figure S8A**), demonstrating that PB2 promotes heterotypic tumor-WBC cluster formation in which tumor cells can possibly evade immune cell killing. To identify which ligand might be involved in CTC-WBC cluster formation, we used a computational ranking of surface proteins to determine which canonical ligands of PB2, i.e., semaphorin interactors^65^, were expressed in WBCs. The online single cell RNA sequencing data suggested that among canonical PB2 ligands, Sema4A had the highest expression within peripheral blood derived mononuclear cells (PBMCs) (macrophages and monocytes), whereas Sema4C was minimally expressed (https://www.proteinatlas.org) (**Figure S8B**). Consistently, the published MS proteomic data^65^ also revealed a higher abundance of Sema4A protein than Sema4D with undetectable Sema4C in human blood monocytes (**Figure 4D**, **Supplementary Excel 1-tab 7**). Furthermore, our flow cytometry analyses of the blood cells from patients with breast cancer demonstrated that, compared to granulocytes and lymphocytes, monocytes express a higher level of Sema4A (**Figure 4E**, **Figure S8C**), and that the heterotypic CTC-WBC clusters double positive for EpCAM and CD45 have enriched co-expression of PB2 and Sema4A versus WBC-only clusters and single cells (**Figure 4F**, **Figure S8D–E**), implying the possible contribution of PB2 and Sema4A to heterotypic CTC-WBC clustering.

To determine the protein interactions between monocyte Sema4A and tumor cell PB2, we utilized the THP1 monocytic cells which express high Sema4A (>80%) (**Figure S8F**) as an alternative source of human monocytes for heterotypic cluster formation. Similar to patient WBCs, they do not impact the viability of PB KO tumor cells in co-culture (**Figure S8G**). Using human tumor cell-THP1 clusters and anti-PB2 for co-immunoprecipitation, we also detected the enrichment of Sema4A and CDC42 in the PB2 protein complex, compared to monocytes only (**Figure 4G**).

To determine the importance of PB2 in tumor-monocyte clustering, we found that in cell suspension co-culture, PB2 control tumor cells formed clusters effectively with THP1 cells at an optimized 1:4 ratio, whereas PB2 KO TNBC cells lost the capability for heterotypic cluster formation (**Figure 4H–I**). Consistently, sorted primary PDX TNBC cells (TN1) with PB2^high^ expression formed significantly larger heterotypic clusters with THP1 cells than PB2^low^ TN1 cells, when co-cultured under the previously established clustering conditions on collagen-coated plates^16^ (**Figure 4J–K**).

To further determine the association of Sema4A with heterotypic CTC clustering, we sorted Sema4A^+^ and Sema4A^-^ monocytes (THP1) for co-culture with PB2 WT and KO TNBC cells (MDA-MB-231). Sema4A^+^ monocytes with PB2^+^ tumor cells (double positive) formed the largest clusters compared to single-negative or double-negative combinations of Sema4A^-/+^ monocytes with PB2^+/-^ tumor cells in heterotypic clustering (**Figure 4L–M**). Interestingly, loss of Sema4A alone was sufficient to reduce the size of heterotypic PB2^+^ tumor clusters to be comparable with PB2 KO cells, suggesting that Sema4A is the primary ligand on monocytes involved in PB2- dependent heterotypic tumor clustering (**Figure 4A**).

### Loss of PB2 from tumor cells reduces metastasis and CTC formation *in vivo*

The existence of CTC clusters in patient blood has previously been shown to be associated with metastasis and reduced OS in breast cancer^11, 22^. Since our work revealed that PB2 is highly expressed on patient CTCs and promotes tumor cell clustering, we continued to determine if PB2 promotes CTC cluster formation and metastasis *in vivo*. We implanted equal numbers of L2G-labeled WT (control) and PB2 KO breast cancer cells orthotopically into the fourth mammary fat pads of NSG mice to monitor spontaneous metastasis to the lungs (**Figure 5A**). Mice were monitored for 10 weeks until the experimental endpoint, at which time they were sacrificed, and tumors, lungs, and blood were collected.

**Figure 5.**
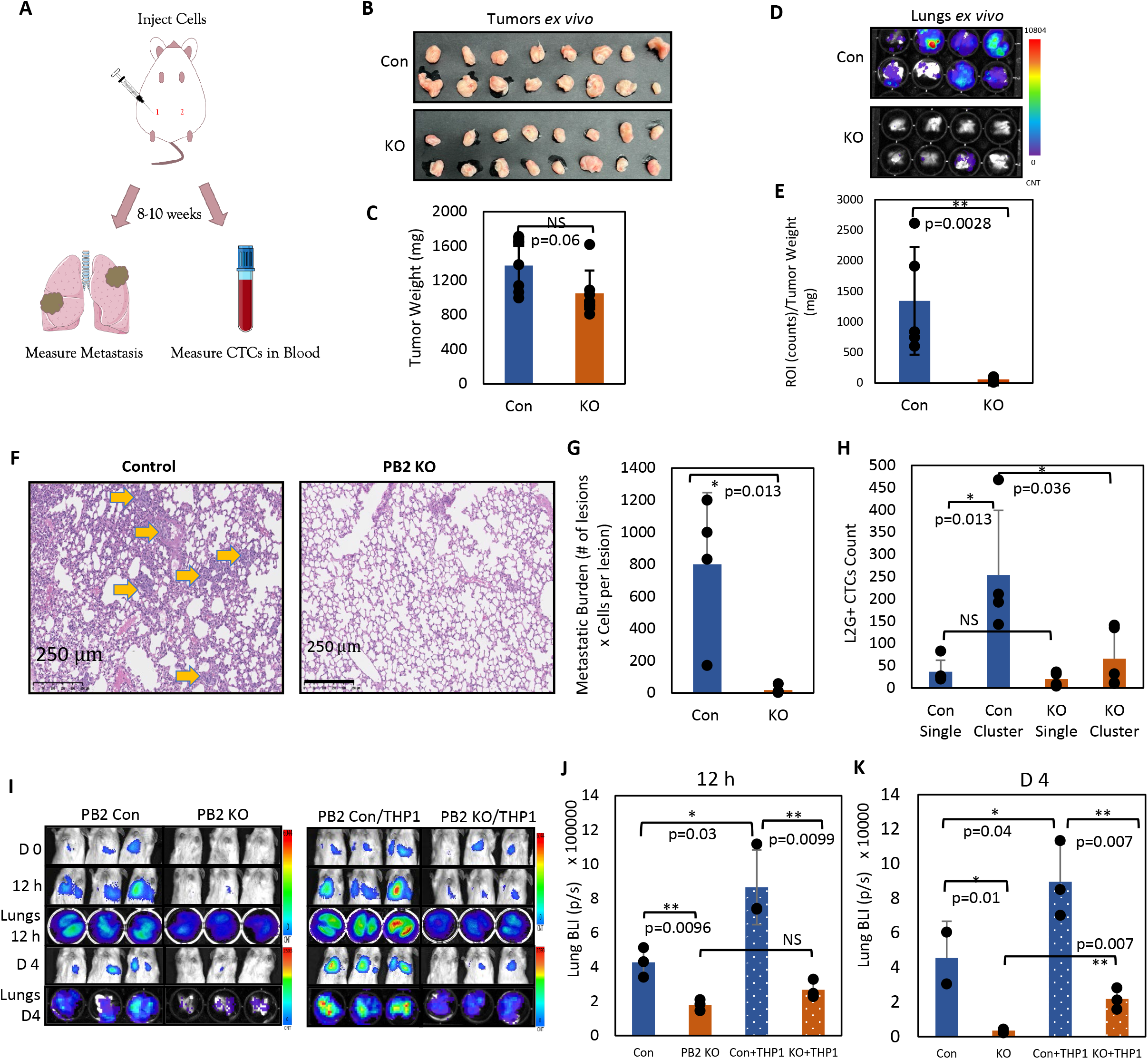
PB2 facilitates metastasis of TNBC *in vivo*. **A)** Schematic showing the experimental workflow of orthotopic implantation of tumor cells into mouse mammary fat pads and subsequent analyses of lung metastases and blood CTCs. **B-C)** Representative images (B) and weight quantification of PB2 Con and KO tumors (C) upon harvest from mice at 10 weeks (N=8). **D-E)** Bioluminescence images of mouse lungs *ex vivo* (D) and normalized lung metastasis burden (E) at 10 weeks, with bioluminescence counts normalized to cumulative tumor weight, N=5 biological replicates. **F-G)** Representative images (F) and metastatic burden (G) in control and PB2 KO metastatic cells of mouse lungs with H&E staining at 10 weeks after orthotopic implantation, scale bar = 250 µm; experiments were repeated with the PB KO2 clone. **H**) L2G^+^ CTC counts (single and clusters) detected in control and PB2 KO mouse blood via flow cytometry at 10 weeks of spontaneous metastasis, N=4. **I**) IVIS bioluminescence images of NSG mice after tail vein injection of pre-clustered MDA- MB-231 cells alone (PB2 Con or KO), and with THP1 cells (PB2 Con/THP1, KO/THP1) on day 0 (D0), 12 h, day 4 (D4). N=3 biological replicates. **J-K)** Mean photon counts/s of ROI (region of interest) of lungs *ex vivo* after harvesting lungs at 12 h (J) and day 4 (K), N=3 biological replicates per time point. Data are presented as mean values +/-SD, with p-values reported from two-sided unpaired t-tests.

Between the control and PB2 KO groups, there was no significant difference in tumor weight (**Figure 5B–C**, **Figure S9A–B**), which was expected given that there is no change in proliferation between PB2 control and KO cells (**Figure S5E**). The lungs, however, showed a significant difference in metastatic burden, with a 23-fold stronger signal from the mice bearing control tumors compared to those of PB2 KO tumors (**Figure 5D–E**). Reduced metastatic colonization (metastatic lesions and cell numbers) in KO tumor-bearing mice was confirmed via H&E staining of lung sections (**Figure 5F–G**). Additionally, blood from each mouse was collected via heart puncture, and L2G^+^ CTCs were analyzed via flow cytometry after red blood cell lysis. There were significantly more cell clusters in the mice with control tumors compared to the mice bearing PB2 KO tumors (**Figure 5H**, **Figure S9C**), demonstrating that PB2 promotes CTC cluster formation and enhances spontaneous metastasis of breast cancer *in vivo*.

To determine if PB2-mediated CTC clustering was responsible for increased metastasis, we compared the outcomes of tail vein-injected PB2 control and KO cancer cells which were pre-clustered for 4 h *ex vivo* prior to injection. PB2 KO tumor cells significantly reduced dissemination to the lungs after 12 h, and the phenotype was maintained for up to four days (**Figure 5I**), suggesting that PB2-mediated clustering specifically promotes metastasis independent of proliferation. Similar results with heterotypic tumor-THP1 clusters were achieved by pre-clustering L2G-labeled tumor cells with unlabeled monocytes. Cells were clustered for 4 h and then injected via tail vein, and L2G^+^ tumor cell dissemination was specifically monitored via bioluminescence imaging at 12 h and 4 days. PB2 Con-THP1 heterotypic clusters mediated significantly more metastases into the lungs compared to PB2 KO-THP1 cells, with the phenotype observed as early at 12 h and holding for up to 4 days (**Figure 5I–K**). Notably, at both 12 h and Day 4 timepoints, the presence of THP1 monocytes in control tumor cell clusters significantly increased the metastatic colonization in the lungs compared to control tumor cell only conditions, whereas THP1 cells had no significant influence on PB2 KO tumor cells (**Figure 5I–K**). Overall, these results show that monocytes promote tumor cell survival and dissemination efficiency to the lungs in a PB2-dependent manner.

## Discussion

Utilizing computational ranking of protein abundance measured by quantitative global proteomics in combination with experimental investigation, our studies identified PB2 as a driver that promotes both homotypic and heterotypic CTC clustering through its interactions with semaphorin ligands Sema4C on tumor cells and Sema4A on monocytes. Additionally, we discovered novel proteins such as MCM7 as downstream mediators of the PB2 interaction network that could facilitate CTC cluster formation and proliferation. Future therapeutics aimed at blocking PB2 and Sema4C/4A interactions can potentially serve as novel targeting strategies in breast cancer especially TNBC, which currently lack target therapeutics.

PB2 has previously been implicated in several cancers, including breast, prostate, and, most notably, brain cancer^51, 59, 60, 66^, partially related to its well-known role in regulating neuronal migration and synaptogenesis^33, 67^. PB2 has been shown to promote invasion in glioblastoma by regulating cellular biomechanics^51, 60^. PB2 also acts as a receptor for angiogenesis, which has been shown to promote stemness in a PB2-dependent manner in brain cancer and prostate cancer^30, 66^. Sema4A is known to be expressed on immune cells and have a role in carcinogenesis^68, 69^. While it is likely that certain cancer types or host immunity may confound the effects of PB2^70^, in the context of advanced breast cancer, PB2 in conjunction with its ligands Sema4C in tumor cells and Sema4A in monocytes drives CTC clustering, metastasis, and is correlated with lower OS and DMFS. However, other known ligands of PB2, such as Sema5A and Sema4D, are either not expressed or not involved in the functions of PB2 in breast cancer. The context-dependent functions and biomarker association of PB2 may be largely dependent on the ligands it interacts with and the microenvironmental conditions^50, 71^. For instance, while Sema4D expression is relatively low in breast cancer cells, its interaction with the ligand of brain endothelial cells may contribute to the brain-tropic metastasis of breast cancer^72^.

Metastasis is a series of complex steps by which tumor cells shed off the primary tumor, enter the vasculature (intravasate), circulate, and ultimately extravasate into secondary tissues^14, 73, 74^. Previous studies demonstrate that myeloid cells such macrophages promote migrating tumor cell contact with the vasculature and enhance vascular permeability for intravasation^75, 76^. Considering the tumor-myeloid interactions promoted by PB2, PB2 may also promote intravasation of tumor cells in addition to CTC clustering and dissemination.

CTC clusters have up to 50-fold greater metastatic potential compared to single CTCs, making them a very crucial population to target. While it has long been hypothesized that CTCs can also cluster with immune cells, limited research has thus far been devoted to identifying the molecular markers of these heterotypic clusters. In addition to myeloid cells such as the monocytes specified in this work, neutrophils, MDMCs^19, 77^, and the plethora of other immune cell types need to be further investigated for potential interactions with and influences on CTCs in cancer metastasis.

## Acknowledgements

This work could not have been completed without the gracious contributions of the core facilities at Northwestern University including the Robert H. Lurie Comprehensive Cancer Center CTC Core, the Flow Cytometry Core, the Pathology Core, the Mouse Histology and Phenotyping Laboratory, and the Center for Comparative Medicine. We are deeply grateful to our collaborators and the Mass Spectrometry Core at Pacific Northwest National Laboratory. We would also like to thank our collaborators and all other Liu laboratory members at Northwestern University Feinberg School of Medicine. This project has been partially supported by Department of Defense grants W81XWH-16-1-0021 and W81XWH-20-1-0679 (H. Liu); NIH/NCI grants R01CA245699 (H.Liu and E.K. Ramos), R01GM139858 (T. Shi), and UG3CA256967 (T. Shi and H. Liu); the Lurie Cancer Center Lynn Sage Breast Cancer Research Foundation (X. Liu, M. Cristofanilli, and H. Liu); American Cancer Society CSCC-Team-23-980420-01-CSCC (H. Liu); a Northwestern University Pharmacology start-up grant (H. Liu); and NIH Fellowships T32 CA009560 (E.K. Ramos), T32 CA080621-15 and the Julius Kahn Fellowship (R. Taftaf), and T32GM008061 (E.J. Schuster).

## Author Contributions

E.S. and N.D. conceived the idea presented here. H.L., T.S, and C-F.T. supervised experimental planning and implementation. N.D. performed initial analysis of survival correlations and protein expression of the candidates in desired breast cancer cell lines. E.S. drove the development of the presented story line and data acquisition. E.S. planned and carried out all cell culture, clustering, imaging, flow cytometry/sorting, immunoprecipitation, and mass spectrometry sample preparation experiments. E.S. planned and completed all animal work *in vivo* with the help of Y.J.. N.D. and K.G. carried out the EV mass spectrometry screen, including all necessary EV isolation and sample preparation. C-F.T. and R.B.K. performed mass spectrometry proteomic analyses of breast cancer cells and the co-immunoprecipitation samples, analyzed and deposited data. C.Z. and R.X. further helped analyze the mass spectrometry proteomic data of EVs. T.Z. and A.H. helped run the computational analysis across multiple cancer data sets to rank surface protein expression in breast cancer. M.C., A.S, W.J.G., L.C.P., C.R. and S.S., and supported blood sample collection from breast cancer patients by enrolling patients, collecting blood work regularly for analysis, and managing patient data. E.S., N.D., Y.J., E.R, R.T., L.E-S., D.S., and V.A.C all helped to process and analyze patient blood samples via flow cytometry. Y.Z. processed and analyzed patient blood samples via CellSearch® with the help of S.S. T.S. and C.F. ran and analyzed multiple mass spectrometry analyses on breast cancer CTCs and clusters. K.P.S. aided in the analysis of TMA data. D.S. preformed ImageStream analysis to validate presence of CTC clusters and immune cells. E.S. and H.L. wrote the manuscript. T.S., Z.T., R.B.K., L.E-S., K.P.S. and V.A.C. edited the text.

## Materials and Methods

### Patient sample collections

Blood samples and tissue sections were collected from stage III-IV breast cancer patients under guidelines from the Institutional Review Board at Northwestern University (IRB protocol STU00203283) in compliance with NIH human subject studies guidelines. Blood samples were collected into CellSave tubes for CellSearch analyses or into EDTA tubes for flow cytometry of live cells. The CellSearch® platform for CTC detection (CD45^-^ DAPI^+^Cytokeratin^+^) had one open immunofluorescence channel for PB2 analysis. Prior to flow cytometry analysis, blood specimens in EDTA tubes underwent red blood cell lysis (Sigma #R7757). Breast tumor tissues and normal adjacent tissues were frozen in tissue freezing buffer OCT until sectioned for laser capture microdissection for mass spectrometry analysis.

### Animal studies

All animal procedures complied with the NIH Guidelines for the Care and Use of Laboratory Animals and were approved by the Northwestern University Institutional Animal Care and Use Committee (IACUC protocol IS00014098). All mice used in this study were kept in specific pathogen-free facilities in the Animal Resources Center at Northwestern University.

### CellSearch

CellSearch analysis processed 7.5 mL of patient blood using the CTC Epithelial Kit (CellSearch #7900001) and CXC Kit (CellSearch #7900017) to deplete immune cells by an EpCAM^+^ selection and identify specific markers. Cells were then stained for CK, DAPI, CD45 (immune cell marker), and PB2 (Miltenyl Biotec #130-126-566).

### Flow cytometry analysis

Mouse/human cells, PDXs, and patient cells were counted and resuspended in wash buffer (PBS+2% FBS). They were blocked with mouse IgG (Sigma #I5381) for 10 minutes on ice and incubated with fluorescence-conjugated antibodies for 15 minutes on ice: PB2 PE (phycoerythrin) (human, R&D Systems #FAB53291P), PB2 APC (human, R&D Systems #FAB53291A), PB2 FITC (mouse, R&D Systems #FAB6836G), CD45 (human, BD Bioscience #557748), EpCAM FITC (human, BD Bioscience #347197), Sema4A PE (human/mouse, BioLegend #148404), Sema4A APC (human/mouse Biolegend #148406). Cells were washed 2x in wash buffer and run for analysis on a fluorescence-associated cell sorting (FACS) LSR cytometer from BD Biosciences.

### Cell sorting

Human/PDX/PBMC cells were resuspended in PBS+2% FBS at a final concentration of 10×10^6^ cells/mL in 5 mL FACS tubes after filtering (Fisher Scientific #352235). Cells were blocked with mouse IgG (Sigma #I5381) for 10 minutes on ice and florescent antibody for 15 minutes on ice: PB2 PE (human, R&D Systems #FAB53291P), PB2 APC (human, R&D Systems #FAB53291A), PB2 FITC (mouse, R&D Systems #FAB6836G), Sema4A PE (human/mouse, BioLegend #148404). Cells were then run through BD FACSMelody Cell Sorter and cells were collected based on gated populations. Collected cells were washed 2x in PBS prior to downstream application.

### Ranking of adhesion proteins based on their abundance

Adhesion proteins (N=608, derived from the Molecular Signature Database^38–40^ of the Gene Ontology Biological Processes^41, 42^) were ranked according to relative protein abundance using proteomics datasets collected from 122 treatment-naive primary breast patient samples^43^, TNBC such as MDA-MB-231 cells^44^, THP1 macrophage cells^78^, and cell-derived EVs. The presence of adhesion proteins of interest in these datasets was first determined. For proteins that were identified in each dataset, they were ranked by spectral counting (the total number of MS/MS spectra for a given protein). Top-ranked proteins from the three published datasets were further cross-examined for downstream analysis.

### Extracellular vesicle isolation

EVs were isolated from human cell lines and PDX models from culture media after 72 h in culture. Media were then pooled and isolated using differential centrifugation. The first centrifugation step was at 2,000 x g for 10 minutes. Media were then transferred into an appropriate ultracentrifuge tube (Beckman Coulter # 344058) and spun at 10,000 x g for 30 minutes. Supernatant was removed and transferred into a clean ultracentrifuge tube and spun at 100,000 x g for 70 minutes. The resulting EV pellet was washed with PBS and spun again at 100,000 x g for 70 minutes. The final EV pellet was resuspended in PBS and stored at -80 °C until analysis.

### Cell culture

Cell lines were maintained in complete supplemented media (10% FBS, 1% penicillin-streptomycin (Sigma-Aldrich P4333-100ML)). MDA-MB-231 and 4T1 cell lines were maintained in DMEM supplemented with high glucose (Corning #10-013-CV) for <20 passages in cell culture incubators at 37 °C, 5% CO2. MDA-MB-468, HS578T, and THP1 cells were cultured in RPMI-1640 supplemented media (Fisher Scientific #SH30027.01) for <20 passages in cell culture incubators at 37 °C, 5% CO2.

### Gene knockdown

A total of 2×10^6^ cells were plated in a 10 cm dishes one day prior to KD (day 0). On the day of KD, ON-TARGETplus siRNA at a final concentration of 50 nM per plate was incubated with Dharmafect 1 at 100 nmol/L (GE Dharmacon #T-2001-03) for 20 minutes at room temperature in reduced serum Opti-MEM Medium (Thermo Fisher Scientific #31985070). siRNA+Dharmafect mixture was added to cells in Opti-MEM for 16-24 h. Cells were passaged in complete media and reseeded at 2×10^6^ cells in a 10 cm dish, and the KD procedure was repeated once more as described. On day 4, cells were harvested, counted, and analyzed via flow cytometry and western blotting to check for sufficient KD of target protein. The following siRNAs were used: SMARTpool Human PLXNB2 (Dharmacon #L-031513-01), SMARTpool Mouse PLXNB2 (Dharmacon #L-040980-00-0010), SMARTpool Human Sema4C (Dharmacon #L-015364-01- 0010), ON-TARGETplus PLXNB2 Human siRNA-09 (Dharmacon #J-031513-09-0010), ON-TARGETplus PLXNB2 Human Individual siRNA-10 (Dharmacon #J-031513-10-0010), ON-TARGETplus PLXNB2 Human Individual siRNA-11 (Dharmacon #J-031513-11-0005), ON-TARGETplus PLXNB2 Human Individual siRNA-12 (Dharmacon #J-031513-12-0005), ON-TARGETplus Non-targeting Pool (Dharmacon #D-001810-10-50).

### Gene overexpression

Cells were transfected with PB2 full-length and mutant overexpression plasmids via a Lipofectamine 3000 Transfection Kit (Thermo Fisher #L3000015). Cells were plated at 300k cells/well in a 6-well plate the day prior to transfection. After transfection, cells were incubated for 48-72 h under standard growth conditions and then harvested to check expression of target protein via flow cytometry and western blotting. The following overexpression vectors were used: pLV-PLXNB2-mRBD (Addgene #86240), pLV-PLXNB2-dVTDL (Addgene 86239), pLV-PLXNB2-dECTO (Addgene #86238), PLXNB2 OHu01778C_pcDNA3.1(+) N-Terminal Flag-Tag (GenScript #SC1626), PLXNB2 OHu01778D_pcDNA3.1+/C- C-Terminal Flag-Tag (GeneScript #OHu01778D), PLXNB2 Untagged Construct (GeneScript #SC1625).

### Gene knockout

Two individual pre-designed human PLXNB2 sgRNA CRISPR-Sanger clones and one non-targeting control vector (Sigma-Aldrich #0020) were ordered from Sigma as glycerol stock (Sigma-Aldrich #HS5000013567, #HS500013568). Bacteria were expanded for maxi prep according to kit protocols (Qiagen #12163). Plasmid was isolated and used to create lentivirus. Cells were infected with either of the sgPLXNB2 clones or the control sgRNA with Cas9-GFP virus (Sigma-Aldrich #0030) at 10 IFU/cell in Opti-MEM (Thermo Fisher Scientific #31985070) with 8 µg/mL supplemented Polybrene (Millipore Sigma #TR-1003-G). Cells were incubated for 4 h at 37 °C in 5% CO2 upon which complete medium was added to the culture. Cells were incubated in normal growth conditions for 48-72 h and monitored for GFP expression. Cells were harvested and analyzed for sufficient KO of protein. KO cells were sorted for multiple pooled clones using the BD FACSMelody Automated Cell Sorter based on PB2 expression.

### Clustering assay

In homotypic tumor cell clustering assays, 20,000 tumor cells were plated in Poly(2-hydroxyethyl methacrylate) (PolyHema)-coated 96-well flat bottom plates (Sigma-Aldrich #P3932-25G). Tumor cells were well mixed to ensure single-cell suspension prior to plating. In heterotypic tumor cell-immune cell clustering assays, tumor cells and immune cells were plated at a 1:4 ratio in PolyHema-coated 96-well flat bottom plates. Cells were monitored in the Incucyte Live Cell Imager (Sartorius) for 24 h and analyzed for average cluster size over time, with a cluster being defined as two or more cells.

### Immunoblotting (western blotting)

Cells were pelleted and then lysed in 1x RIPA Lysis Buffer (VWR Amresco #N653-100mL) supplemented with 1:100 protease inhibitor cocktail (Thermo Fisher #78440) and incubated on ice for 30 minutes. Lysates were then centrifuged for 10 minutes at 14,600 x g at 4 °C. Cell lysates were measured for protein concentration using Bradford analysis (Thermo Fisher #23209, Bio-Rad #500-0006). Totals of 10-30 µg of protein lysate and 10 µL of dual color protein standard ladder (Bio-Rad #161-0374) were loaded onto 4-20% Mini-PROTEAN gels (Bio-Rad #4568094) and then transferred to nitrocellulose membranes using the Bio-Rad TurboTransfer system and kits (Bio-Rad #1704270). Membranes were blocked with 5% milk in Tris-buffered saline and 0.1% Tween 20 (TBST) for 60 minutes and then washed in 0.1% TBST. Primary antibody was incubated on membranes in 5% milk in TBST for 1.5 h at room temperature or at 4 °C overnight. Secondary HRP-conjugated antibodies were added at a dilution of 1:10,000 in 5% milk in TBST and incubated for 60 minutes at room temperature (Anti-Mouse: Promega #W402B; Anti-Rabbit: Promega #W401B). The Pierce SuperSignal West Pico PLUS chemiluminescent substrate (Thermo Scientific #34577) was added one minute prior to imaging using the Bio-Rad Chemidoc imaging system. Primary antibodies used include: Anti-PB2 (Protein Tech #10602-1-AP), Anti-Bactin (Abcam #AB8224), Anti-Sema4A (Thermo Fisher #PA5- 101258), Anti-Sema4C (Ray Biotech #102-11819), Anti-Sema4G (Santa Cruz Biotech #sc- 515644), Anti-Sema4D/CD100 (Santa Cruz Biotech #sc-39065), Anti-Sema7C (Santa Cruz Biotech #sc-376149).

### Co-immunoprecipitation

The co-immunoprecipitation protocol was done as directed in the Dynabeads Co-Immunoprecipitation Kit (Thermo Scientific #14321D). PB2 (Protein Tech #10602-1-AP) or control rabbit IgG (Protein Tech #3000-0-AP) were pre-conjugated to Dynabeads at 7 µg antibody/mL of beads. Cells were lysed using immunoprecipitation-lysis buffer and incubated overnight at 4 °C with 7.5 mg of pre-conjugated Dynabeads for every 1-15 g protein. Beads were washed and protein was eluted using SDS loading buffer for downstream applications of western blotting and mass spectrometry analysis.

### Cell preparation for mass spectrometry analysis

MDA-MB-231 breast cancer tumor cells underwent double transfection of siPB2 for KD of PB2 (Dharmacon #L-031513-01) and control siRNAs (Dharmacon #D-001810-10-50). Cells were then trypsinized and resuspended on PolyHema-coated plates and allowed to cluster for 4 h. Adherent cells were scraped from the plate at the zero hour time point. After clustering, cells were harvested and lysed using RIPA buffer (VWR Amresco #N653-100mL) supplemented 1:100 with protease inhibitor (Thermo Fisher #78440).

### Protein digestion by S-Trap for proteomic analysis

MDA-MB-231 cell lysates were resuspended in SDS buffer (the final concentration was 5%). The protein concentration was measured with a BCA protein assay (Thermo Fisher Scientific). A total of 50 mg of protein was reduced with 10 mM DTT for 15 minutes at 37 °C and subsequently alkylated with 50 mM iodoacetamide at 25 °C for 15 minutes in the dark. The samples were acidified by phosphoric acid (the final concentration was 2.5%) and then diluted with six volumes of “binding” buffer (90% methanol; 100 mM triethylammonium bicarbonate, TEAB; pH 7.1). After mixing, the protein solution was loaded onto an S-Trap filter from Protifi (Huntington, NY), spun at 10,000 g for 1 minute, and then the filter was washed with 150 μL of binding buffer three times. Proteins were digested with Lys-C (Wako) and sequencing-grade trypsin (Promega, V5117) (1 μg of each in 20 μL of 50 mM TEAB) in the filter at 37 °C for 16 h. To elute peptides, 40 μL of 50 mM TEAB, 40 μL of 0.2% formic acid (FA) in H2O, and 40 μL of 50% acetonitrile (CAN) in H2O were added sequentially. The peptide solutions were pooled for BCA assay to estimate peptide amounts. Totals of 20 μg of peptides were dried with a SpeedVac and stored at -80 °C until LC-MS/MS analysis.

### LC-MS/MS analysis

Lyophilized peptides were reconstituted in 200 μL of 0.1% TFA with 2% ACN containing 0.01% n-Dodecyl β-D-maltoside (DDM)^79^ to reach a concentration of 0.1 μg/μL, and 5 μL of the resulting sample was analyzed by LC-MS/MS using an Orbitrap Fusion Lumos Tribrid mass spectrometer (Thermo Scientific) connected to a nanoACQUITY UPLC system (Waters) (buffer A: 0.1% FA with 3% ACN and buffer B: 0.1% FA in 90% ACN) as previously described^80^. Peptides were separated by a gradient mixture with an analytical column (75 μm i.d. × 20 cm) packed using 1.9-μm ReproSil C18 and with a column heater set at 50 °C using an LC gradient of 2-6% buffer B in 1 min, 6-30% buffer B in 84 min, 30-60% buffer B in 9 min, 60-90% buffer B in 1 min, and finally 90% buffer B for 5 min at 200 nL/min. The DIA-MS/MS scan was performed in the HCD mode with the following parameters: precursor ions from 350–1650 *m/z* were scanned at 120,000 resolution with an ion injection time of 60 ms and an AGC target of 1E6. The range of *m/z* (isolation window) of data-independent acquisition (DIA) windows from 377 (54), 419 (32), 448 (28), 473.5 (25), 497.5 (25), 520.5 (23), 542.5 (23), 564.5 (23), 587 (24), 610.5 (25), 635 (26), 660 (26), 685.5 (27), 712.5 (29), 741 (30), 771 (32), 803.5 (35), 838.5 (37), 877 (42), 921 (48), 972 (52), 1034.5 (71), 1133.5 (129), and 1423.5 (453) was scanned at 30,000 resolution with an ion injection time of 120 ms and an AGC target of 3E6. The isolated ions were fragmented with HCD at the 30% level.

### Proteomic data analysis

The raw DIA data were processed with Fragpipe^81, 82^ and searched against a human UniProt database (FASTA file dated Jan. 05, 2022 with 40,818 sequences which contained 20,409 decoys). Initial fragment mass tolerances were set to 20 ppm. A peptide search was performed with strict tryptic digestion (Trypsin) and allowed a maximum of two missed cleavages. Carbamidomethyl (C) was set as a fixed modification; acetylation (protein N-term) and oxidation (M) were set as variable modifications. DIA_SpecLib_Quant workflow was used for DIA quantitation (takes DIA data as input, builds a spectral library using MSFragger-DIA, then quantifies with DIA-NN).

### PDX mouse models

Multiple PDX models of human breast cancer were previously established in the lab^13, 16^. Cells from TNBC patient tumors or pleural effusion were used to establish tumors that propagated in immunodeficient NSG mice. PDX models were labeled with either Luc2- tdTomato (L2T) (red) or Luc2-eGFP (L2G) (green) reporters to track tumor growth and measure metastasis using bioluminescence imaging and fluorescence analyses (microscopy or flow cytometry). Models were maintained in the lab through tumor harvesting, dissociation, and re- implantation of tumor cells into the mammary fat pads of NSG mice. PDX models were routinely checked for expression/lack of expression of key markers to monitor phenotype. PDX models were sorted for PB2^high/low^ cells for downstream *in vitro* experiments and analysis.

### Lung colonization *via* tail vein injection and bioluminescence imaging

All mice used in this study were female NSG mice 6-12 weeks of age (Jackson Laboratory) and housed in the pathogen-free barrier facility in the Animal Resources Facility at Northwestern University. All animal studies were completed under approved protocols (IS00014098) and adhered to all procedures and regulations outlined by the NIH Guidelines for the Care and Use of Laboratory Animals. Animals were randomized by age and weight and excluded from experiments if presenting conditions unrelated to tumors. Mice were injected with 200,000 L2G or L2T-labeled tumor cells via the tail vein, and lung colonization was monitored upon intraperitoneal injection of luciferin (Gold Bio #LUCK-1G 115144-35-9). The bioluminescence signals of metastasis burden were imaged with the IVIS Spectrum Imager, using same imaging time (acquisition times ranged from 5 s to 5 min) across all groups with identical region of interest quantified for comparison. Mice were kept four days post-injection or until the survival endpoint. At the experimental endpoint, lungs were harvested, imaged in PBS *ex vivo* for bioluminescence on black-wall 24 well plates, and then preserved in formalin (Fisher Scientific #SF98-4) for H& E staining.

### Spontaneous metastasis *in vivo*

Mice were injected with 10,000 L2T/L2G-labeled tumor cells in each lower mammary fat pad. Tumors were monitored weekly for growth, and lung metastasis was monitored using bioluminescence imaging. Tumors grew for 8 weeks or until survival endpoint, at which point tumors, lungs, and blood were collected and analyzed *ex vivo.* Lungs were imaged under bioluminescence and microscopy and preserved in formalin (Fisher Scientific #SF98-4) for hematoxylin-eosin (H&E) staining and immunohistochemistry (IHC) analysis. Tumors were weighed and preserved in formalin for H&E/IHC analysis. Blood was collected, and red blood cells were lysed using RBC lysis buffer (Sigma #R7757) followed by analysis via flow cytometry for L2G/L2T-positive tumor cells.

### Tumor microarray (TMA) and IHC

A total of 89 formalin-fixed paraffin-embedded breast tumor tissues were included in the tumor TMA, with selected tumor regions guided by H&E-stained images. To make a TMA that allows microscopic comparison of the staining characteristics of different blocks while preventing exhaustion of pathological material, a core of paraffin was removed from a “recipient” paraffin block (one without embedded tissue), and the remaining empty space was filled with a core of paraffin-embedded tissue from a “donor” block. An H&E-stained recipient block that is representative of the tissue remaining in the donor block was used to select the sample core with a color marker corresponding to tumor, benign, etc. Matched blocks were pulled out and a recipient TMA block was made and trimmed with the face of the block even with the size of a 1.5 mm core by using the semi-automatic Veridiam Tissue Microarrayer VTA-100. The created TMA block was sectioned for staining. In this TMA, 19 cases from the STU00203283, 9 ER-negative cases, 30 triple-negative cases, 27 ER-positive cases, and 4 normal breast cases (see **Supplementary Excel 3**) were selected and used to construct the recipient block. Detailed patient information can be found in supplementary table. IHC was performed with the help of Bella Shmaltsuyeva of the Robert H. Lurie Comprehensive Cancer Center Pathology Core Facility with PB2 antibody (Protein Tech #10602-1-AP).

### H&E staining

Mouse lungs and tumors were paraffin-embedded, and sections were mounted on slides and processed with H&E staining with help from the Mouse Histology and Phenotyping Laboratory at Northwestern University.

### Mammosphere assay

Tumor cells were plated at 1,000 cells per well on PolyHema-coated 12- well plates in Prime-XV Tumorsphere SFM media (Irvine Scientific #91130). Cells were monitored for up to 10 days and the total number of mammospheres per well was counted. Mammospheres were defined as groups with 25 or more cells originating from a single cell.

### Quantification and statistical analysis

All data unless otherwise specified is displayed as mean ± standard deviation (SD). Statistical analysis was done using Microsoft Excel to calculate p-values using two-tailed Student’s t-tests, with p£0.05 considered significant.

## Data Availability

The RAW global MS data and the identified results from Fragpipe and DIA-NN have been deposited in Japan ProteOme STandard Repository (jPOST: https://repository.jpostdb.org/). The accession codes: JPST002098 for jPOST and PXD041009 for ProteomeXchange. The access link is JPST002098 for jPOST and P PXD041009 for ProteomeXchange. The access link is https://repository.jpostdb.org/preview/20324360326419dd4f77ff8 and access key is 3715.

## Supplementary Figure Legends

**Figure S1.**
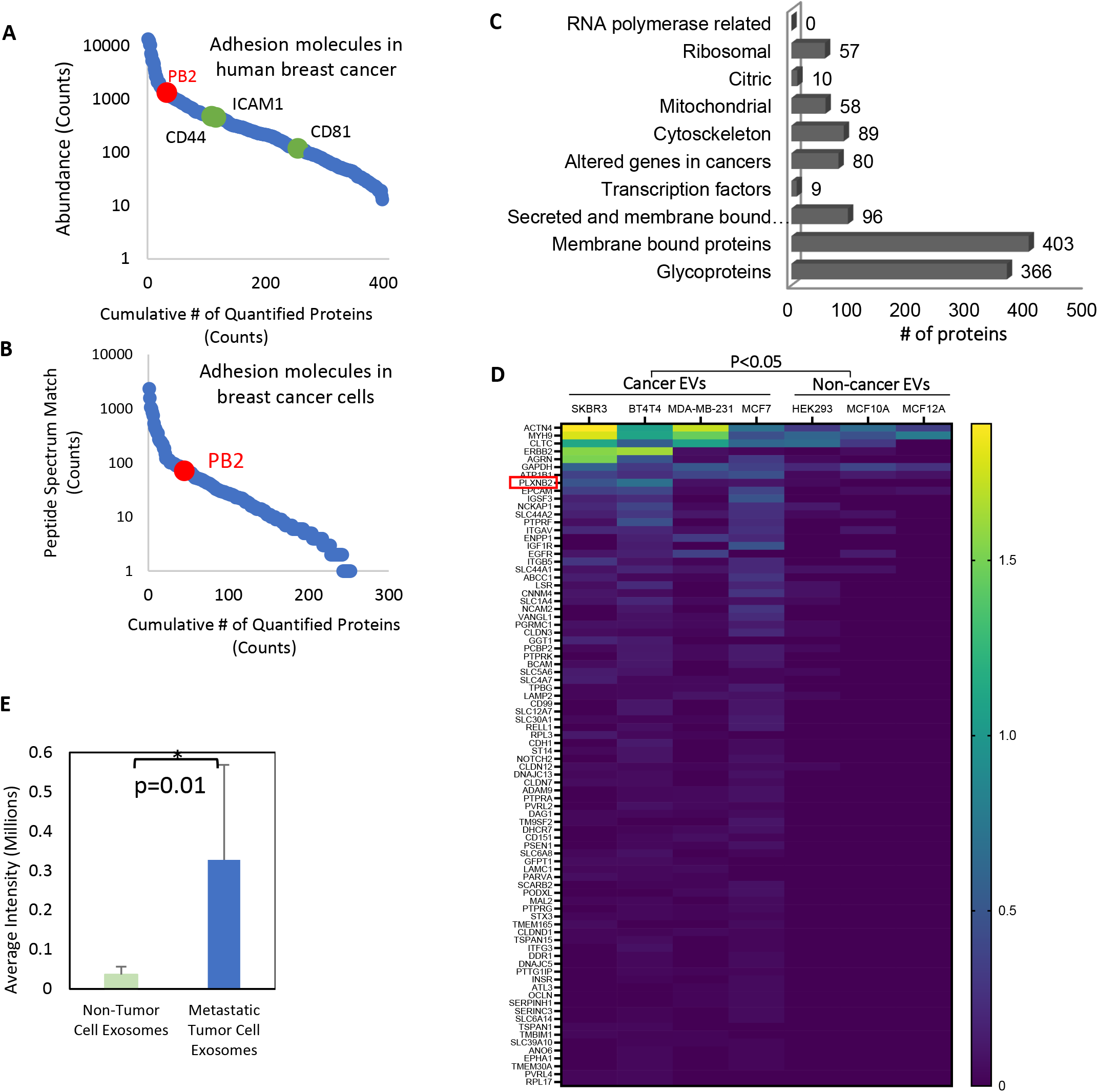
PB2 as a top candidate in primary breast tumors. **A**) Average spectral counts-based abundance ranking of 398 identified adhesion/surface proteins across 122 treatment-naïve primary breast patient samples (N=122) (https://doi.org/10.1016/j.cell.2020.10.036), highlighting key proteins previously identified to be significant in CTCs (CD44, ICAM1, CD81 in green) and a novel candidate PB2 at a higher rank (red). **B**) Average peptide spectrum match (PSM)-based abundance ranking of 252 identified adhesion/surface proteins in MDA-MB-231 triple negative breast cancer (TNBC) cell lines (N=3) (https://doi.org/10.1021/acs.jproteome.1c00293). **C**) Categories of MS-identified extracellular vesicle proteins (total 1603 peptides) secreted by normal and cancer cells. **D**) Heat map of MS-identified differentially expressed proteins between EVs derived from breast cancer cell lines vs. EVs from immortalized normal epithelial cells (unpaired t test P<0.05). PB2 is indicated by a red box. **E**) Table showing the top 11 proteins significantly up-regulated in breast tumor tissue vs. normal tissue and whether or not they significantly correlate to poor prognosis in breast cancer. **F**) Average intensity expression (in millions) of PB2 in EVs of normal non-tumor epithelial cells compared to that of metastatic breast cancer cells. Data are reported as mean +/- SD, p-value is from a two-sided unpaired t-test.

**Figure S2.**
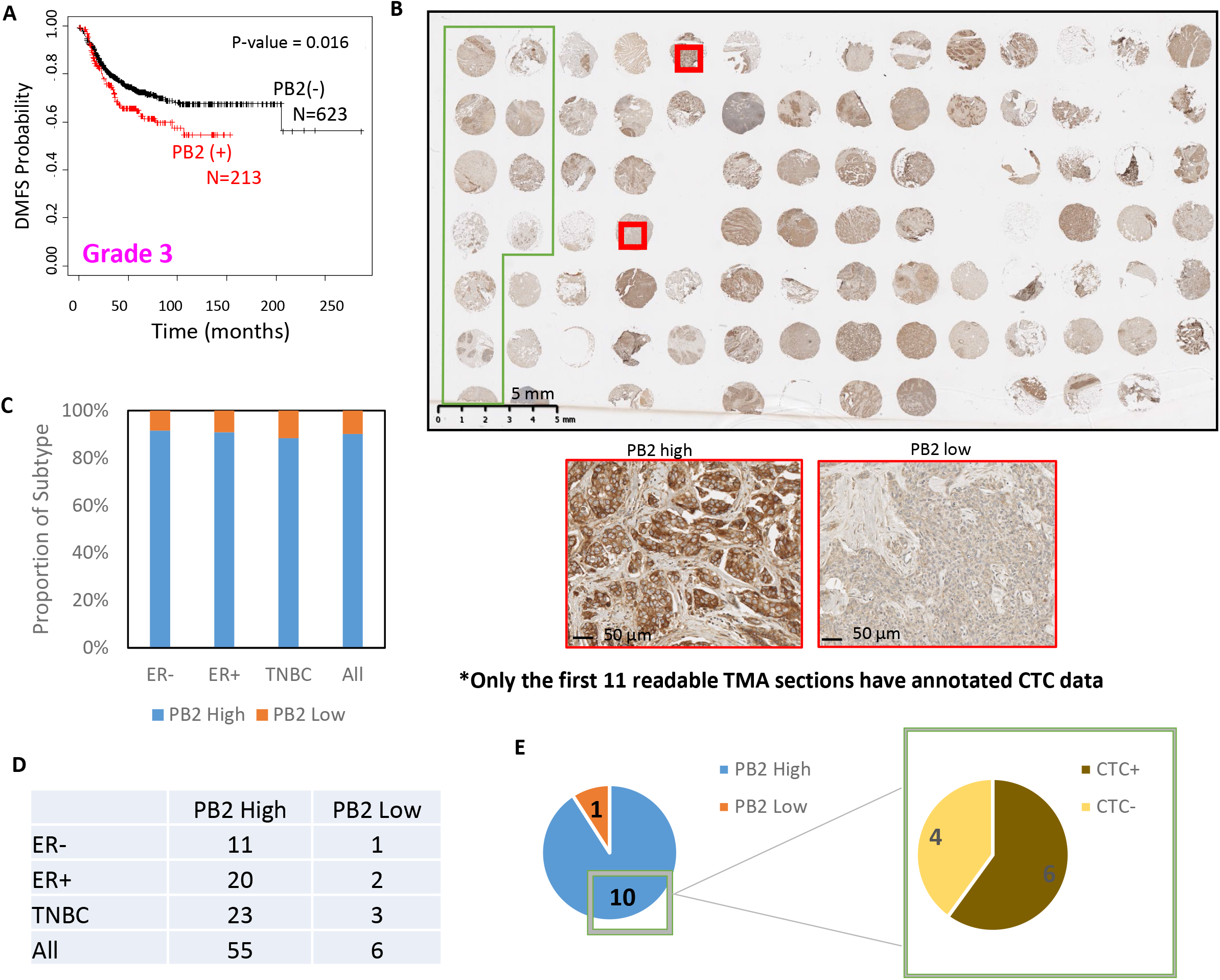
PB2 expression in human breast tumors is associated with detection of CTCs. **A**) KM plot for distant metastasis-free survival (DMFS) of grade 3 breast cancer with high vs. low PB2 mRNA expression, N=836 (data from KMPlotter). **B**) Tissue microarray of advanced breast cancer patient samples subjected to IHC staining for PB2 (N=86). The first 11 cases from the B06 cohort of patients (STU00203283), as highlighted by the green box on the TMA (see Suppl. Excel 3), had simultaneous analyses for the number of CTCs in the blood. **C & D)** Annotation of PB2 status of TMA patients stratified by specific molecular subtype of breast cancer (C) and tabular annotation of the cases (D). **E**) PB2 status of first 11 patients from the B06 cohort in correlation with CTC status in blood.

**Figure S3.**
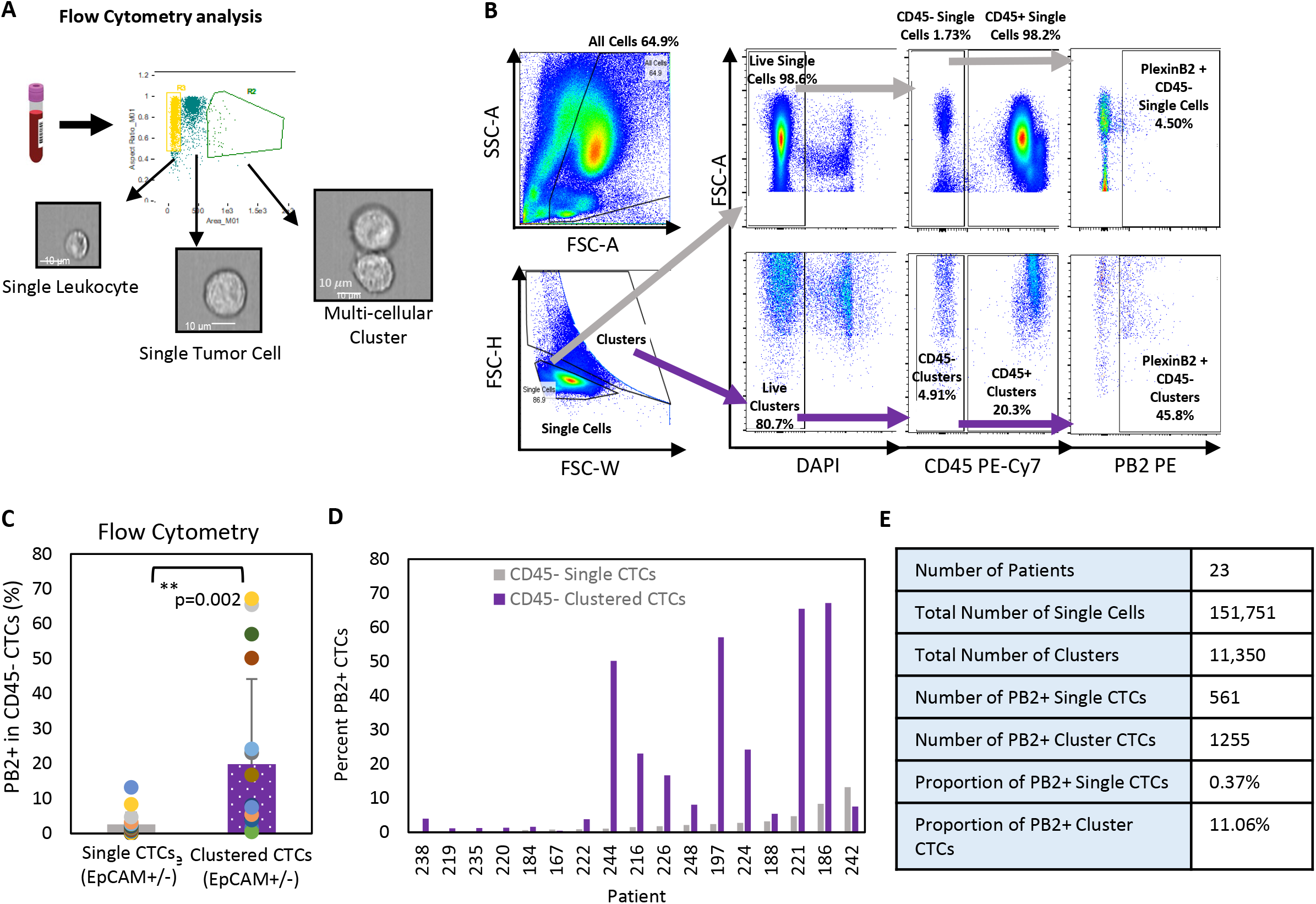
PB2 expression analysis in patient CTCs via flow cytometry. **A**) Schematic showing workflow for flow cytometry analysis of blood cells from patients with breast cancer. **B**) Representative flow cytometry gating of single and clustered blood cells, by PB2^+/-^ and CD45^+/-^ status, from the B06 cohort of advanced breast cancer patients. **C**) Average percentage of cells expressing PB2 in CD45^-^ single and clustered CTCs from flow analysis, including all EpCAM^+/-^ CTCs, N=17 patients, data reported as mean values +/- SD, p-value from a two-sided unpaired t-test. **D**) Percentage of PB2^+^ CD45^-^ single and CD45^-^ clustered CTCs from advanced breast cancer patients at the time of blood draw, N=17 patients, data stratified by individual patient. **E**) Table showing total number and proportion of CTC single cells or CTC clusters that are PB2^+^ from breast cancer patients as reported by flow analysis, N=23.

**Figure S4.**
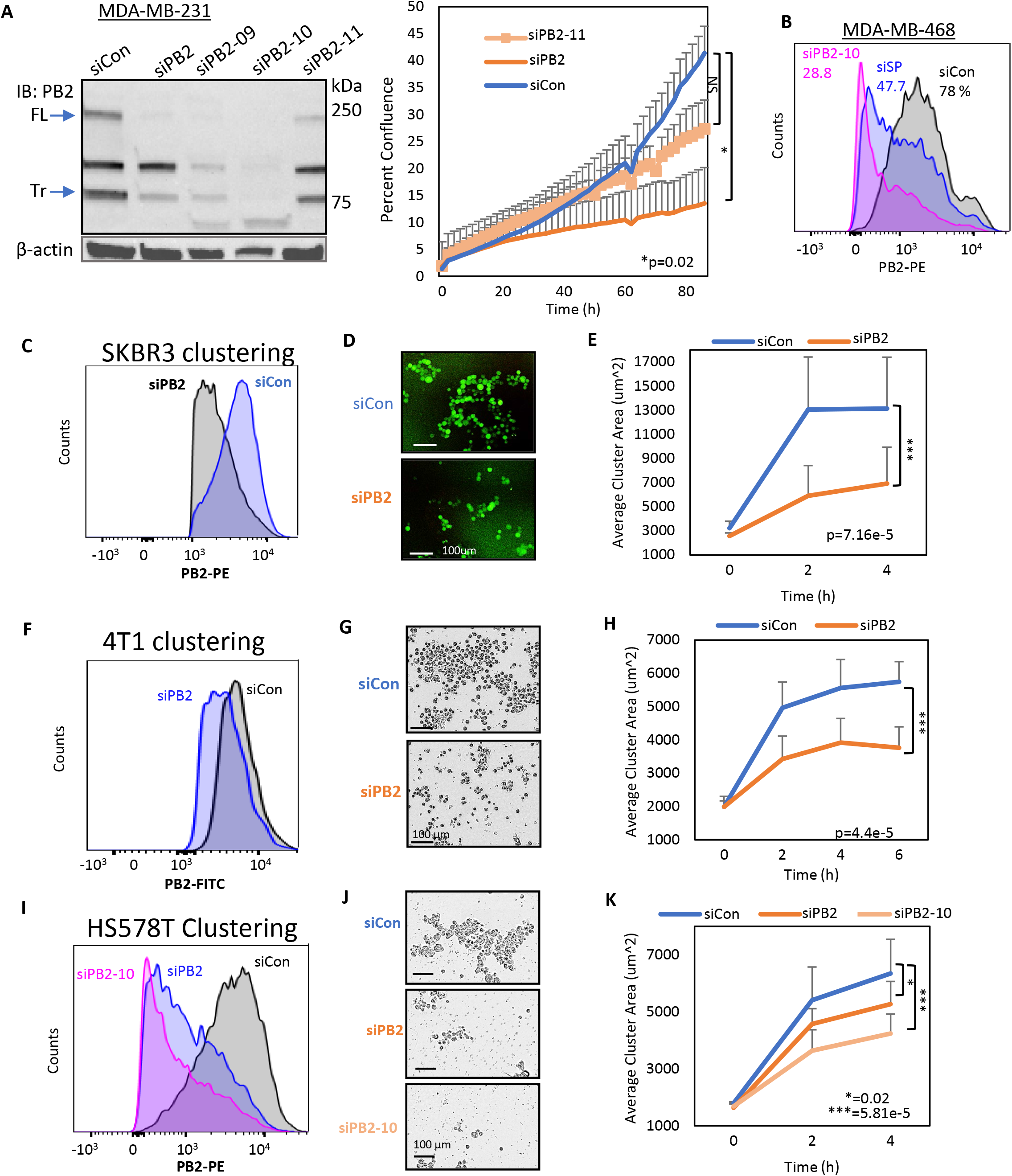
PB2 knockdown inhibits clustering of human and mouse breast cancer cells. **A**) **Left panel:** Western blot showing MDA-MB-231 KD efficiency of PB2 using SmartPool (siPB2) siRNA as well as single siRNAs (si09, si10, si11), N=3 experiments; **Right panel:** Proliferation of MDA-MB-231 cells measured as percent confluence using Incucyte. Breast cancer cells plated after double transfection with either SmartPool siRNA (siPB2) or single siRNA KD of PB2 (siPB2-11), N=3 biological replicates with at least 3 technical replicates each, data are presented as mean +/- SD, p-value at 84 h reported as two-sided unpaired t-test. **B**) Flow analysis showing MDA-MB-468 KD of PB2 using SmartPool (siPB2) siRNA as well as single siRNA (siPB2-10), N=3 experiments. **C**) Flow panel showing KD efficiency of SKBR3 breast cancer cells after double transfection with PB2 SmartPool siRNA. **D**) Representative images of SKBR3 cell clusters after 4 h as taken by the Incucyte Live Cell Imager. **E**) Quantification of clusters of SKBR3 cells with PB2 KD after 4 h, average cluster area measured by IncuCyte, with N=4 experiments. Data are presented as mean values +/- SD, p-value is from a two-sided unpaired t-test. **F**) Flow panel showing KD efficiency of 4T1 mouse breast cancer cells after double transfection with PB2 SmartPool siRNA. **G**) Representative images of 4T1 cell clusters after 6 h as taken by the Incucyte Live Cell Imager. **H**) Quantification of clustering assay of 4T1 cells with PB2 KD after 6 h, average cluster area measured by Incucyte, with N=3 experiments. Data are presented as mean values +/- SD, p-value is from a two-sided unpaired t-test. **I**) Flow panel showing KD efficiency of HS578T breast cancer cells after double transfection with PB2 SmartPool siRNA and siPB2-10 single PB2 siRNA. **J**) Representative images of HS578T cell clusters after 4 h as taken by the Incucyte Live Cell Imager. **K**) Quantification of clustering assay of HS578T cells with PB2 KD after 4 h, average cluster area measured by Incucyte, with N=3 experiments. Data are presented as mean values +/- SD, p-value is from a two-sided unpaired t-test.

**Figure S5.**
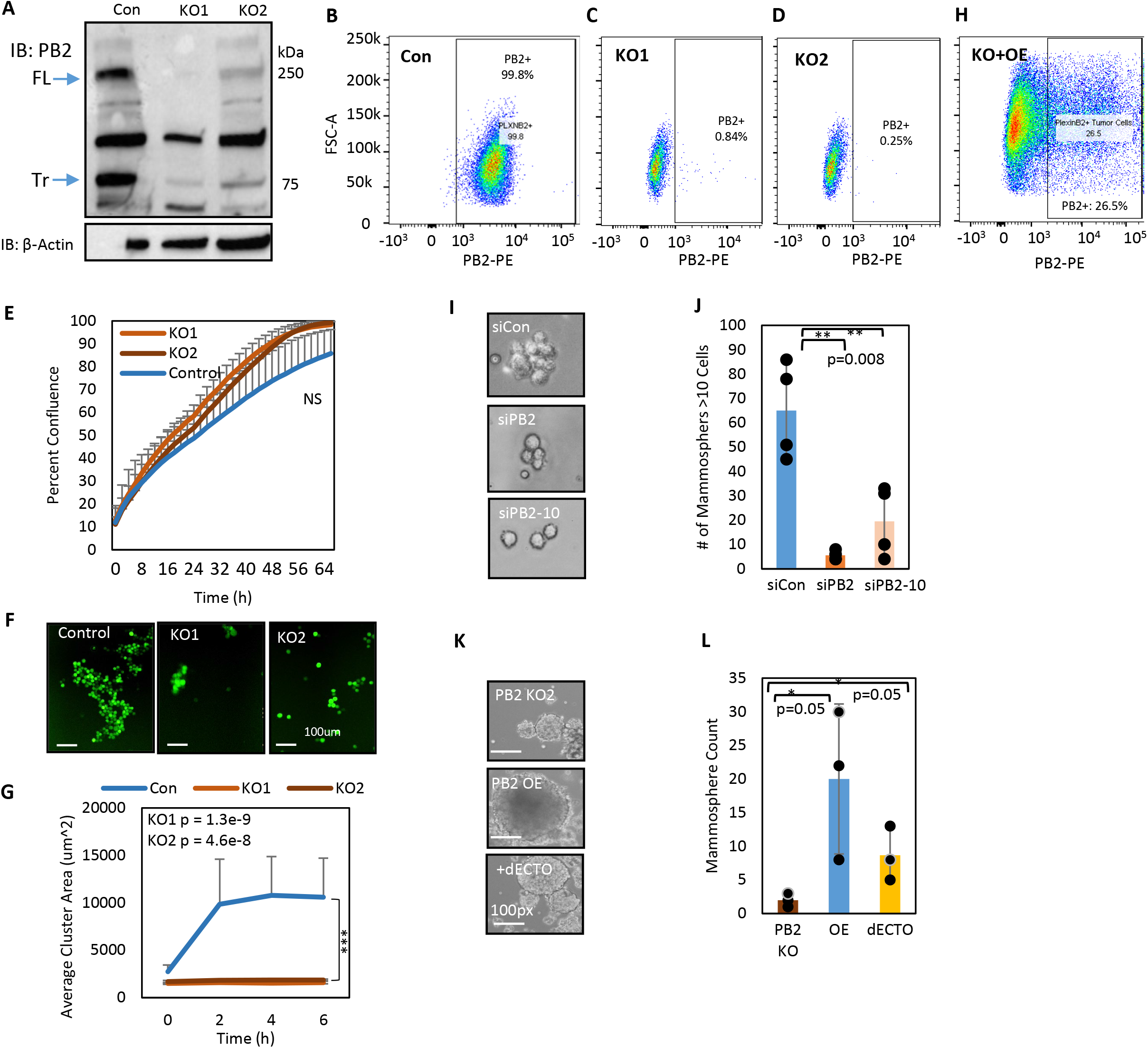
PB2 KO inhibits clustering and mammosphere formation. **A**) Immunoblots of full-length (FL) and truncated (Tr) PB2 in MDA-MB-231 cells with two pooled clones of CRISPR/Cas9-mediated KO, KO1 and KO2, with PB2-specific bands at 250 kDa and 75 kDa, N=3 experiments; **B-D)** Flow cytometry analysis of MDA-MB-231 Cas9 control cells for PB2, KO1, and KO2. **E)** Proliferation assay of MDA-MB-231 control and KO cells over 60+ h measured as percent confluence by IncuCyte, N=3 experiments, data reported as mean +/- SD, p-value is from a two-sided unpaired t-test. **F-G)** Representative images (F) and cluster size quantification (G) of MDA-MB-231 PB2 control, KO1, and KO2 cell clusters after 6 h as taken by the IncuCyte Live Cell Imager; N=3 experiments with at least 3 technical replicates each, data are presented as mean values +/- SD, p-value is from a two-sided unpaired t-test. **H**) Flow cytometry analysis of PB2 overexpression in MDA-MB-231 PB2 KO cells. **I & J)** Representative images (I) of average mammosphere counts of 10 cells or more (J) of MDA-MB-231 WT cells with scramble (siCon), PB2 SmartPool (siPB2), and PB2 single (siPB2-10) siRNA KD after 7 days; N=3 experiments with at least 5 technical replicates each, data are presented as mean values +/- SD, p-values are from a two-sided unpaired t-test. **K & L)** Mammosphere images (K) and counts (L) of MDA-MB-231 KO2 cells with overexpression of PB2 full-length (PB2 OE) and a PB2 mutant with depleted extracellular domain (dECTO) for 10 days, N=4 replicates (bottom). Mammospheres larger than 100 microns across were counted. Data are presented as mean +/-SD, p-value is from a two-sided unpaired t-test.

**Figure S6.**
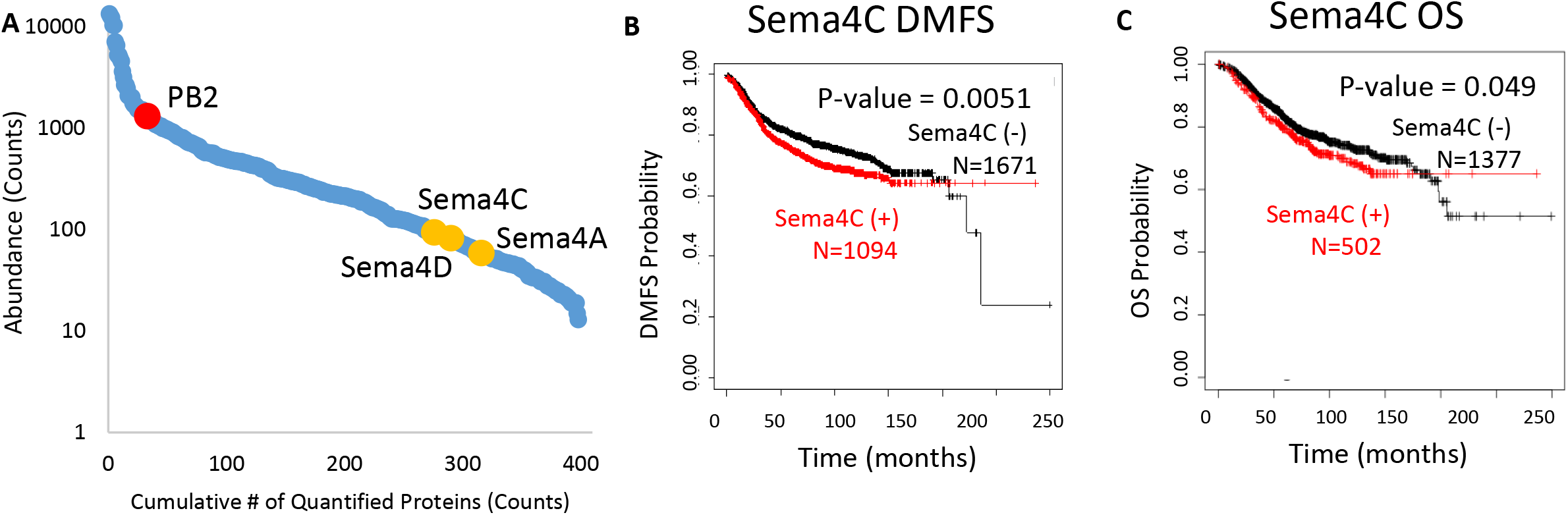
Sema4C ranking and association with patient outcomes. **A**) Abundance ranking of PB2 and Sema4 family members based on average spectral counts among 398 identified adhesion/surface proteins across 122 treatment-naïve primary breast patient samples (https://doi.org/10.1016/j.cell.2020.10.036); the canonical binding ligands of PB2, Sema4C, Sema4D, and Sema4A, are shown by yellow dots. **B**) KM plot of distant metastasis-free survival (DMFS) of breast cancer patients based on mRNA expression of Sema4C, p=0.0051. **C**) KM plot of overall survival (OS) of breast cancer patients based on mRNA expression of Sema4C, p=0.049.

**Figure S7.**
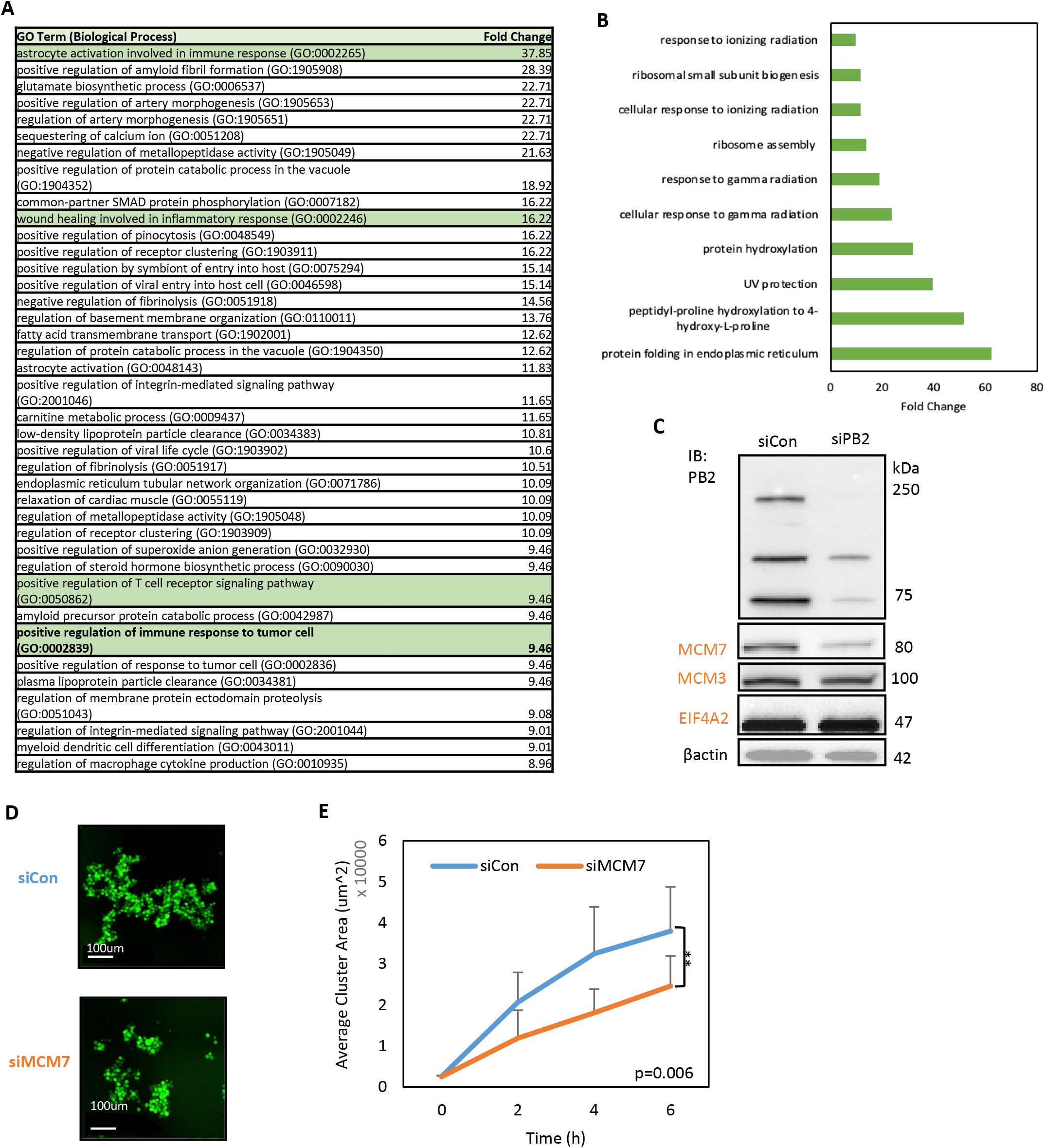
PB2-regulated pathways and downstream target MCM7. **A**) Table showing fold change in the top hits from GO analysis of biological processes for proteins that were significantly up-regulated in PB2 KD compared to control cells by global mass spectrometry analysis of clustered cells. **B**) GO biological processes analysis of altered proteins in Groups C and F in Figure 3H, with clustering-specific up-regulation and down-regulation, respectively, and reversed in a siPB2- dependent manner. **C**) Western blot validation of top hits from Group B (see Figure 3H) of global mass spectrometry analysis of siPB2 single cells vs. clusters. **D & E)** Representative images (D) and average cluster area (E) of MDA-MB-231 control vs. MCM7 KD cells as taken by the IncuCyte. N=3 experiments with at least 3 technical replicates each, data are presented as mean values +/- SD, p-value is from a two-sided unpaired t-test.

**Figure S8.**
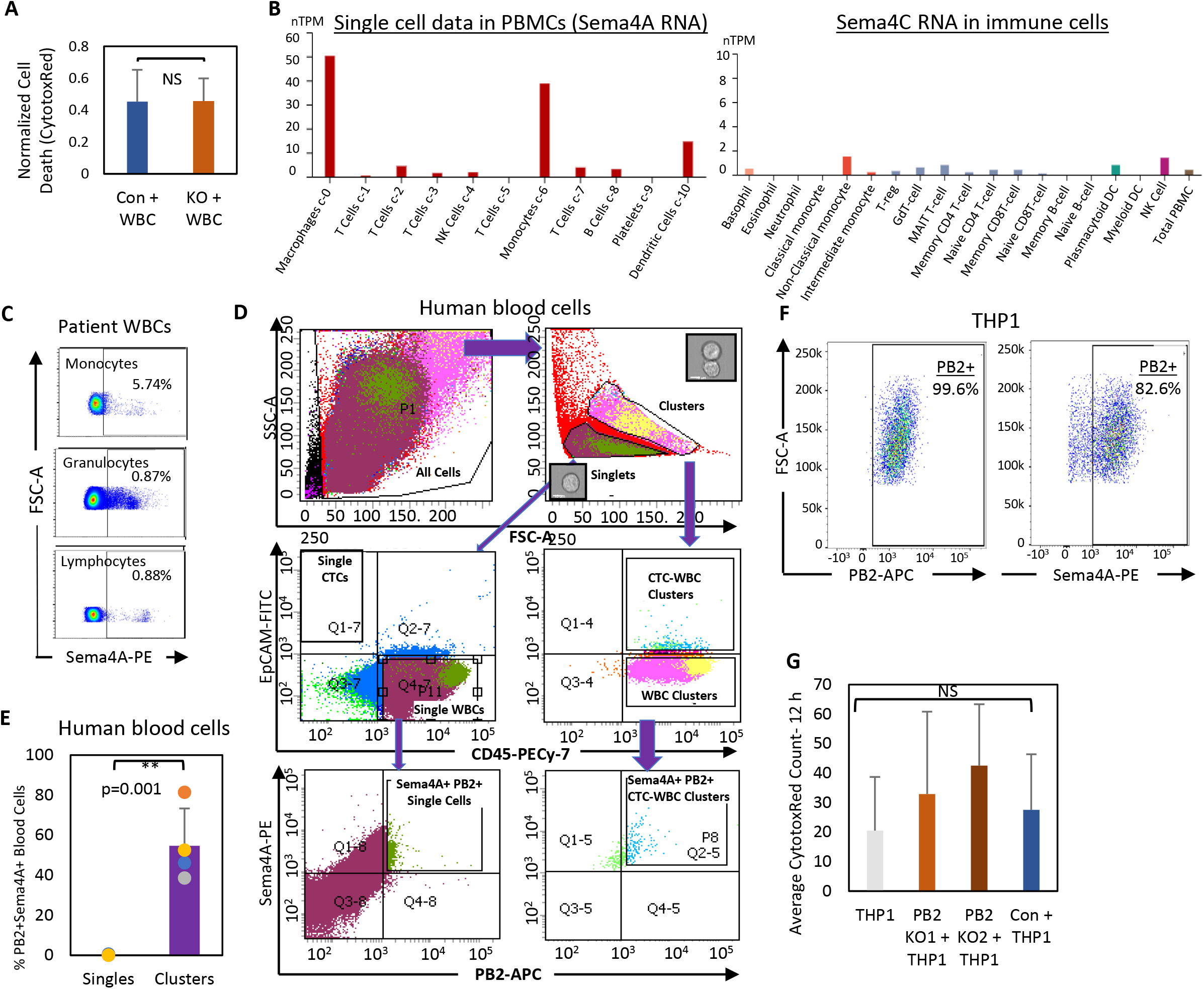
Sema4A expression in human WBCs and monocytes. **A**) Quantification of normalized cytotoxic red dye-labeled dead cell events as measured by IncuCyte at 12 h during the MDA-MB-231 breast cancer cell co-culture with WBCs isolated from breast cancer patients, N=4. Data reported as mean +/-SD, p-value is from a two-sided unpaired t-test. **B**) Sema4A expression in human PBMCs, analyzed via single-cell sequencing of the Human Protein Atlas. **C**) Representative flow plots of Sema4A expression in monocytes, granulocytes, and lymphocytes derived from the blood of patients with breast cancer. **D**) Representative gating strategy of flow cytometry analysis of advanced stage breast cancer patient WBCs stained for EpCAM, CD45, PB2, and Sema4A to identify PB2^+^Sema4A^+^ heterotypic CTC clusters. **E**) Quantification of double positive PB2^+^Sema4A^+^ clusters (CD45^+/-^) in advanced stage breast cancer PBMCs, N=4 patients, data are presented as mean +/-SD, p-value is from a two-sided unpaired t-test. **F**) Flow cytometry analysis of THP1 monocyte cell line for PB2 and Sema4A expression, N=3. **G**) Average Cytotox Red counts in heterotypic THP1 co-culture system as counted by IncuCyte, N=3, data are presented as mean +/-SD, p-value is from an unpaired two-sided t-test.

**Figure S9.**
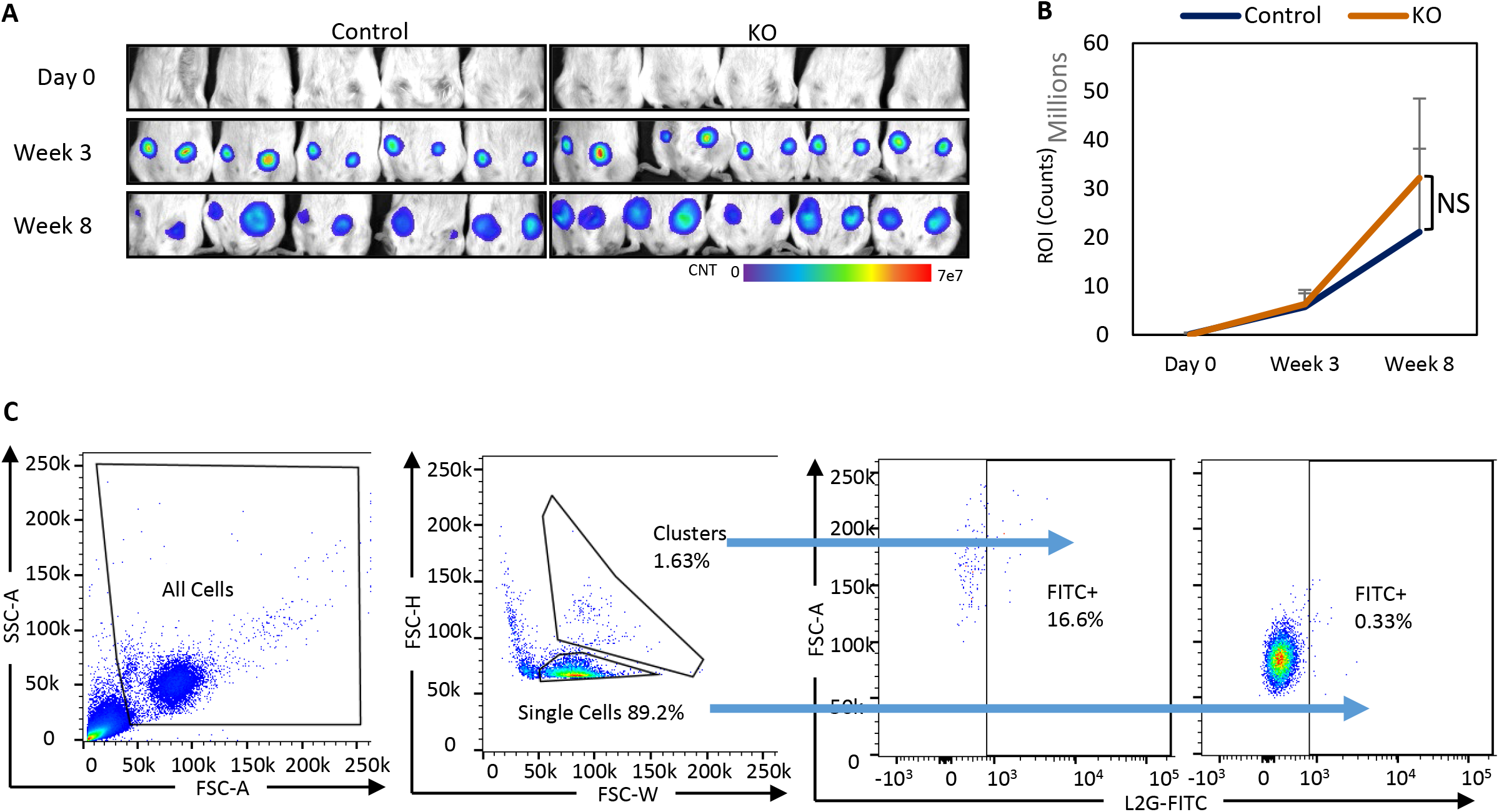
PB2 KO inhibits CTC cluster formation in spontaneous metastasis of TNBC. **A**) Tumorigenesis of MDA-MB-231 control and KO1 tumor cells labeled with L2G luciferase reporter in NSG mice over 8 weeks as shown by bioluminescence imaging, starting with implantation of 10,000 cells per injection side, two injections per mice, N=5 biological replicates. **B**) Average ROI counts of bioluminescence imaging from tumorigenesis (A) as measured by an SII Lago imager, data reported as mean +/-SD over time, N=5 biological replicates, p-value from an unpaired two-sided t-test. **C**) Representative flow analysis of L2G^+^ CTCs from mouse blood following 10 weeks of spontaneous metastasis, N=5.

## References

1 Min, S., Lee, B. & Yoon, S. Deep learning in bioinformatics. Brief Bioinform 18, 851–869, doi:10.1093/bib/bbw068 (2017).

2 Miotto, R., Wang, F., Wang, S., Jiang, X. & Dudley, J. T. Deep learning for healthcare: review, opportunities and challenges. Brief Bioinform 19, 1236–1246, doi:10.1093/bib/bbx044 (2018).

3 Xu, J. et al. Interpretable deep learning translation of GWAS and multi-omics findings to identify pathobiology and drug repurposing in Alzheimer’s disease. Cell Rep 41, 111717, doi:10.1016/j.celrep.2022.111717 (2022).

4 Stahlschmidt, S. R., Ulfenborg, B. & Synnergren, J. Multimodal deep learning for biomedical data fusion: a review. Brief Bioinform 23, doi:10.1093/bib/bbab569 (2022).

5 Kang, M., Ko, E. & Mersha, T. B. A roadmap for multi-omics data integration using deep learning. Brief Bioinform 23, doi:10.1093/bib/bbab454 (2022).

6 DeSantis, C. E. et al. Breast cancer statistics, 2019. CA Cancer J Clin 69, 438–451, doi:10.3322/caac.21583 (2019).

7 Loibl, S., Poortmans, P., Morrow, M., Denkert, C. & Curigliano, G. Breast cancer. Lancet 397, 1750–1769, doi:10.1016/S0140-6736(20)32381-3 (2021).

8 Takada, M. & Toi, M. Neoadjuvant treatment for HER2-positive breast cancer. Chin Clin Oncol 9, 32, doi:10.21037/cco-20-123 (2020).

9 Li, X. et al. Triple-negative breast cancer has worse overall survival and cause-specific survival than non-triple-negative breast cancer. Breast Cancer Res Treat 161, 279–287, doi:10.1007/s10549-016-4059-6 (2017).

10 Chen, W., Hoffmann, A. D., Liu, H. & Liu, X. Organotropism: new insights into molecular mechanisms of breast cancer metastasis. NPJ Precis Oncol 2, 4, doi:10.1038/s41698-018-0047-0 (2018).

11 Aceto, N. et al. Circulating tumor cell clusters are oligoclonal precursors of breast cancer metastasis. Cell 158, 1110–1122, doi:10.1016/j.cell.2014.07.013 (2014).

12 Dashzeveg, N. K. et al. New Advances and Challenges of Targeting Cancer Stem Cells. Cancer Res 77, 5222–5227, doi:10.1158/0008-5472.CAN-17-0054 (2017).

13 Liu, H. et al. Cancer stem cells from human breast tumors are involved in spontaneous metastases in orthotopic mouse models. Proc Natl Acad Sci U S A 107, 18115–18120, doi:1006732107 [pii] 10.1073/pnas.1006732107 (2010).

14 Massague, J. & Obenauf, A. C. Metastatic colonization by circulating tumour cells. Nature 529, 298–306, doi:10.1038/nature17038 (2016).

15 Cristofanilli, M. et al. The clinical use of circulating tumor cells (CTCs) enumeration for staging of metastatic breast cancer (MBC): International expert consensus paper. Crit Rev Oncol Hematol 134, 39–45, doi:10.1016/j.critrevonc.2018.12.004 (2019).

16 Liu, X. et al. Homophilic CD44 Interactions Mediate Tumor Cell Aggregation and Polyclonal Metastasis in Patient-Derived Breast Cancer Models. Cancer Discov 9, 96–113, doi:10.1158/2159-8290.CD-18-0065 (2019).

17 Pineiro, R., Martinez-Pena, I. & Lopez-Lopez, R. Relevance of CTC Clusters in Breast Cancer Metastasis. Adv Exp Med Biol 1220, 93–115, doi:10.1007/978-3-030-35805-1_7 (2020).

18 Schuster, E. et al. Better together: circulating tumor cell clustering in metastatic cancer. Trends Cancer 7, 1020–1032, doi:10.1016/j.trecan.2021.07.001 (2021).

19 Szczerba, B. M. et al. Neutrophils escort circulating tumour cells to enable cell cycle progression. Nature 566, 553–557, doi:10.1038/s41586-019-0915-y (2019).

20 Clotilde Costa, L. M.-R., Victor Cebey-López, Thais Pereira-Veiga, Inés Martínez-Pena, Manuel Abreu, Alicia Abalo, Ramón M Lago-Lestón, Carmen Abuín, Patricia Palacios, Juan Cueva, Roberto Piñeiro, Rafael López-López. Analysis of a Real-World Cohort of Metastatic Breast Cancer Patients Shows Circulating Tumor Cell Clusters (CTC-clusters) as Predictors of Patient Outcomes. Cancers 12, doi:10.3390/cancers12051111 (2020).

21 Jansson, S., Bendahl, P. O., Larsson, A. M., Aaltonen, K. E. & Ryden, L. Prognostic impact of circulating tumor cell apoptosis and clusters in serial blood samples from patients with metastatic breast cancer in a prospective observational cohort. BMC Cancer 16, 433, doi:10.1186/s12885-016-2406-y (2016).

22 Mu, Z. et al. Prospective assessment of the prognostic value of circulating tumor cells and their clusters in patients with advanced-stage breast cancer. Breast Cancer Res Treat 154, 563–571, doi:10.1007/s10549-015-3636-4 (2015).

23 Murlidhar, V. et al. Poor Prognosis Indicated by Venous Circulating Tumor Cell Clusters in Early-Stage Lung Cancers. Cancer Res 77, 5194–5206, doi:10.1158/0008-5472.CAN-16-2072 (2017).

24 Wang, C. et al. Longitudinally collected CTCs and CTC-clusters and clinical outcomes of metastatic breast cancer. Breast Cancer Res Treat 161, 83–94, doi:10.1007/s10549-016-4026-2 (2017).

25 Taftaf, R. et al. ICAM1 initiates CTC cluster formation and trans-endothelial migration in lung metastasis of breast cancer. Nat Commun 12, 4867, doi:10.1038/s41467-021-25189-z (2021).

26 Ramos, E. K. et al. Machine learning-assisted elucidation of CD81-CD44 interactions in promoting cancer stemness and extracellular vesicle integrity. Elife 11, doi:10.7554/eLife.82669 (2022).

27 Gkountela, S. et al. Circulating Tumor Cell Clustering Shapes DNA Methylation to Enable Metastasis Seeding. Cell 176, 98–112 e114, doi:10.1016/j.cell.2018.11.046 (2019).

28 Hurtado, P., Martinez-Pena, I., & Piñerio, R. . Dangerous Liasons: Circulating Tumor Cells (CTCs) and Cancer-Associated Fibroblasts (CAFs). Cancers 12, doi:10.3390/cancers12102861 (2020).

29 Cheung, K. J. et al. Polyclonal breast cancer metastases arise from collective dissemination of keratin 14-expressing tumor cell clusters. Proc Natl Acad Sci U S A 113, E854–863, doi:10.1073/pnas.1508541113 (2016).

30 Yu, W. et al. Plexin-B2 Mediates Physiologic and Pathologic Functions of Angiogenin. Cell 171, 849–864 e825, doi:10.1016/j.cell.2017.10.005 (2017).

31 Zielonka, M., Xia, J., Friedel, R. H., Offermanns, S. & Worzfeld, T. A systematic expression analysis implicates Plexin-B2 and its ligand Sema4C in the regulation of the vascular and endocrine system. Exp Cell Res 316, 2477–2486, doi:10.1016/j.yexcr.2010.05.007 (2010).

32 Atkin-Smith, G. K. et al. Plexin B2 Is a Regulator of Monocyte Apoptotic Cell Disassembly. Cell Rep 29, 1821–1831 e1823, doi:10.1016/j.celrep.2019.10.014 (2019).

33 Deng, S. et al. Plexin-B2, but not Plexin-B1, critically modulates neuronal migration and patterning of the developing nervous system in vivo. J Neurosci 27, 6333–6347, doi:10.1523/JNEUROSCI.5381-06.2007 (2007).

34 Junqueira Alves, C. et al. Plexin-B2 orchestrates collective stem cell dynamics via actomyosin contractility, cytoskeletal tension and adhesion. Nat Commun 12, 6019, doi:10.1038/s41467-021-26296-7 (2021).

35 Li, Y. et al. Macrophages facilitate peripheral nerve regeneration by organizing regeneration tracks through Plexin-B2. Genes Dev 36, 133–148, doi:10.1101/gad.349063.121 (2022).

36 Zhou, X. et al. Microglia and macrophages promote corralling, wound compaction and recovery after spinal cord injury via Plexin-B2. Nat Neurosci 23, 337–350, doi:10.1038/s41593-020-0597-7 (2020).

37 Maier, V. et al. Semaphorin 4C and 4G are ligands of Plexin-B2 required in cerebellar development. Mol Cell Neurosci 46, 419–431, doi:10.1016/j.mcn.2010.11.005 (2011).

38 Subramanian, A. et al. Gene set enrichment analysis: a knowledge-based approach for interpreting genome-wide expression profiles. Proc Natl Acad Sci U S A 102, 15545–15550, doi:10.1073/pnas.0506580102 (2005).

39 Liberzon, A. et al. Molecular signatures database (MSigDB) 3.0. Bioinformatics 27, 1739–1740, doi:10.1093/bioinformatics/btr260 (2011).

40 Liberzon, A. et al. The Molecular Signatures Database (MSigDB) hallmark gene set collection. Cell Syst 1, 417–425, doi:10.1016/j.cels.2015.12.004 (2015).

41 Ashburner, M. et al. Gene ontology: tool for the unification of biology. The Gene Ontology Consortium. Nat Genet 25, 25–29, doi:10.1038/75556 (2000).

42 Gene Ontology, C. The Gene Ontology resource: enriching a GOld mine. Nucleic Acids Res 49, D325–D334, doi:10.1093/nar/gkaa1113 (2021).

43 Krug, K. et al. Proteogenomic Landscape of Breast Cancer Tumorigenesis and Targeted Therapy. Cell 183, 1436–1456 e1431, doi:10.1016/j.cell.2020.10.036 (2020).

44 Kim, H. et al. Quantitative Proteomics Reveals Knockdown of CD44 Promotes Proliferation and Migration in Claudin-Low MDA-MB-231 and Hs 578T Breast Cancer Cell Lines. J Proteome Res 20, 3720–3733, doi:10.1021/acs.jproteome.1c00293 (2021).

45 Chen, I. H. et al. Phosphoproteins in extracellular vesicles as candidate markers for breast cancer. Proc Natl Acad Sci U S A 114, 3175–3180, doi:10.1073/pnas.1618088114 (2017).

46 Dai, J. et al. Exosomes: key players in cancer and potential therapeutic strategy. Signal Transduct Target Ther 5, 145, doi:10.1038/s41392-020-00261-0 (2020).

47 Sandfeld-Paulsen, B. et al. Exosomal Proteins as Diagnostic Biomarkers in Lung Cancer. J Thorac Oncol 11, 1701–1710, doi:10.1016/j.jtho.2016.05.034 (2016).

48 Zhou, B. et al. Application of exosomes as liquid biopsy in clinical diagnosis. Signal Transduct Target Ther 5, 144, doi:10.1038/s41392-020-00258-9 (2020).

49 Lanczky, A. & Gyorffy, B. Web-Based Survival Analysis Tool Tailored for Medical Research (KMplot): Development and Implementation. J Med Internet Res 23, e27633, doi:10.2196/27633 (2021).

50 Uhlen, M. et al. A pathology atlas of the human cancer transcriptome. Science 357, doi:10.1126/science.aan2507 (2017).

51 Huang, Y. et al. Plexin-B2 facilitates glioblastoma infiltration by modulating cell biomechanics. Commun Biol 4, 145, doi:10.1038/s42003-021-01667-4 (2021).

52 Charafe-Jauffret, E. et al. Breast cancer cell lines contain functional cancer stem cells with metastatic capacity and a distinct molecular signature. Cancer Res 69, 1302–1313, doi:10.1158/0008-5472.CAN-08-2741 (2009).

53 Grillet, F. et al. Circulating tumour cells from patients with colorectal cancer have cancer stem cell hallmarks in ex vivo culture. Gut 66, 1802–1810, doi:10.1136/gutjnl-2016-311447 (2017).

54 Yang, M. H., Imrali, A. & Heeschen, C. Circulating cancer stem cells: the importance to select. Chin J Cancer Res 27, 437–449, doi:10.3978/j.issn.1000-9604.2015.04.08 (2015).

55 Malik, M. F., Ye, L. & Jiang, W. G. The Plexin-B family and its role in cancer progression. Histol Histopathol 29, 151–165, doi:10.14670/HH-29.151 (2014).

56 Worzfeld, T. & Offermanns, S. Semaphorins and plexins as therapeutic targets. Nat Rev Drug Discov 13, 603–621, doi:10.1038/nrd4337 (2014).

57 Worzfeld, T. et al. Genetic dissection of plexin signaling in vivo. Proc Natl Acad Sci U S A 111, 2194–2199, doi:10.1073/pnas.1308418111 (2014).

58 Conrotto, P., Corso, S., Gamberini, S., Comoglio, P. M. & Giordano, S. Interplay between scatter factor receptors and B plexins controls invasive growth. Oncogene 23, 5131–5137, doi:10.1038/sj.onc.1207650 (2004).

59 Gurrapu, S., Pupo, E., Franzolin, G., Lanzetti, L. & Tamagnone, L. Sema4C/PlexinB2 signaling controls breast cancer cell growth, hormonal dependence and tumorigenic potential. Cell Death Differ 25, 1259–1275, doi:10.1038/s41418-018-0097-4 (2018).

60 Le, A. P. et al. Plexin-B2 promotes invasive growth of malignant glioma. Oncotarget 6, 7293–7304, doi:10.18632/oncotarget.3421 (2015).

61 Xia, J. et al. Semaphorin-Plexin Signaling Controls Mitotic Spindle Orientation during Epithelial Morphogenesis and Repair. Dev Cell 33, 299–313, doi:10.1016/j.devcel.2015.02.001 (2015).

62 Bray, K. et al. Cdc42 overexpression induces hyperbranching in the developing mammary gland by enhancing cell migration. Breast Cancer Res 15, R91, doi:10.1186/bcr3487 (2013).

63 Qu, K. et al. MCM7 promotes cancer progression through cyclin D1-dependent signaling and serves as a prognostic marker for patients with hepatocellular carcinoma. Cell Death Dis 8, e2603, doi:10.1038/cddis.2016.352 (2017).

64 Wei, C. et al. Crosstalk between cancer cells and tumor associated macrophages is required for mesenchymal circulating tumor cell-mediated colorectal cancer metastasis. Mol Cancer 18, 64, doi:10.1186/s12943-019-0976-4 (2019).

65 Ravenhill, B. J., Soday, L., Houghton, J., Antrobus, R. & Weekes, M. P. Comprehensive cell surface proteomics defines markers of classical, intermediate and non-classical monocytes. Sci Rep 10, 4560, doi:10.1038/s41598-020-61356-w (2020).

66 Li, S., Goncalves, K. A., Lyu, B., Yuan, L. & Hu, G. F. Chemosensitization of prostate cancer stem cells in mice by angiogenin and plexin-B2 inhibitors. Commun Biol 3, 26, doi:10.1038/s42003-020-0750-6 (2020).

67 Duan, Y. et al. Semaphorin 5A inhibits synaptogenesis in early postnatal- and adult-born hippocampal dentate granule cells. Elife 3, doi:10.7554/eLife.04390 (2014).

68 Ito, D. & Kumanogoh, A. The role of Sema4A in angiogenesis, immune responses, carcinogenesis, and retinal systems. Cell Adh Migr 10, 692–699, doi:10.1080/19336918.2016.1215785 (2016).

69 Liu, X. et al. Sema4A Responds to Hypoxia and Is Involved in Breast Cancer Progression. Biol Pharm Bull 41, 1791–1796, doi:10.1248/bpb.b18-00423 (2018).

70 Karki, A. et al. Plexin-B2 and Semaphorins Do Not Drive Rhabdomyosarcoma Proliferation or Migration. Sarcoma 2022, 9646909, doi:10.1155/2022/9646909 (2022).

71 Uhlen, M. et al. Proteomics. Tissue-based map of the human proteome. Science 347, 1260419, doi:10.1126/science.1260419 (2015).

72 Klotz, R. et al. Circulating Tumor Cells Exhibit Metastatic Tropism and Reveal Brain Metastasis Drivers. Cancer Discov 10, 86–103, doi:10.1158/2159-8290.CD-19-0384 (2020).

73 Aceto, N., Toner, M., Maheswaran, S. & Haber, D. A. En Route to Metastasis: Circulating Tumor Cell Clusters and Epithelial-to-Mesenchymal Transition. Trends Cancer 1, 44–52, doi:10.1016/j.trecan.2015.07.006 (2015).

74 Yu, M. et al. Circulating breast tumor cells exhibit dynamic changes in epithelial and mesenchymal composition. Science 339, 580–584, doi:10.1126/science.1228522 (2013).

75 Roh-Johnson, M. et al. Macrophage contact induces RhoA GTPase signaling to trigger tumor cell intravasation. Oncogene 33, 4203–4212, doi:10.1038/onc.2013.377 (2014).

76 Harney, A. S. et al. Real-Time Imaging Reveals Local, Transient Vascular Permeability, and Tumor Cell Intravasation Stimulated by TIE2hi Macrophage-Derived VEGFA. Cancer Discov 5, 932–943, doi:10.1158/2159-8290.CD-15-0012 (2015).

77 Sprouse, M. L. et al. PMN-MDSCs Enhance CTC Metastatic Properties through Reciprocal Interactions via ROS/Notch/Nodal Signaling. Int J Mol Sci 20, doi:10.3390/ijms20081916 (2019).

78 Zhang, T. et al. A proteome-wide assessment of the oxidative stress paradigm for metal and metal-oxide nanomaterials in human macrophages. NanoImpact 17, doi:10.1016/j.impact.2019.100194 (2020).

79 Tsai, C. F. et al. Surfactant-assisted one-pot sample preparation for label-free single-cell proteomics. Commun Biol 4, 265, doi:10.1038/s42003-021-01797-9 (2021).

80 Tsai, C. F. et al. An Improved Boosting to Amplify Signal with Isobaric Labeling (iBASIL) Strategy for Precise Quantitative Single-cell Proteomics. Mol Cell Proteomics 19, 828–838, doi:10.1074/mcp.RA119.001857 (2020).

81 Teo, G. C., Polasky, D. A., Yu, F. & Nesvizhskii, A. I. Fast Deisotoping Algorithm and Its Implementation in the MSFragger Search Engine. J Proteome Res 20, 498–505, doi:10.1021/acs.jproteome.0c00544 (2021).

82 Demichev, V. et al. dia-PASEF data analysis using FragPipe and DIA-NN for deep proteomics of low sample amounts. Nat Commun 13, 3944, doi:10.1038/s41467-022-31492-0 (2022).

